# Natural genetic variation determines microglia heterogeneity in wild-derived mouse models of Alzheimer’s disease

**DOI:** 10.1101/2020.06.02.130237

**Authors:** Hongtian Stanley Yang, Kristen D. Onos, Kwangbom Choi, Kelly J. Keezer, Daniel A. Skelly, Gregory W. Carter, Gareth R. Howell

**Author notes:** Submission corresponding authors and. equal contributing authors (Hongtian Stanley Yang, Kristen D. Onos).

## Abstract

Microglia are now considered drivers of Alzheimer’s disease (AD) pathology. However, single-cell RNA-sequencing (scRNA-seq) of microglia in mice, a key preclinical model organsim, have shown mixed results regarding translatability to human studies. To address this, scRNA-seq of microglia from C57BL/6J (B6) and wild-derived strains WSB/EiJ, CAST/EiJ and PWK/PhJ carrying *APP/PS1* was performed and demonstrated that genetic diversity significantly altered features and dynamics of microglia in baseline neuroimmune functions and in response to amyloidosis. There was significant variation in abundance of microglial subpopulations, including numbers of disease-associated microglia and interferon-responding microglia across the strains. Further, for each subpopulation, significant gene expression differences were observed between strains, and relative to B6 that included nineteen genes previously associated with human AD including *Apoe, Trem2, Bin1* and *Sorl1*. This resource will be critical in the development of appropriately targeted therapeutics for AD and a range of other neurological diseases.

## Introduction

Alzheimer’s disease (AD) is defined by the neuropathological accumulation of beta amyloid plaques, neurofibrillary tangles of tau and widespread neuronal loss. AD is the most common cause of adult dementia and is characterized by a wide range of cognitive and behavioral deficits that severely impact quality of life and the ability to self-care. Recent work has re-focused the field towards the contribution of brain glial cells to the initiation and spread of these disease-specific pathologies, and specifically on the role of microglia as potentially a causative cell type in driving disease development and progression. Human genome-wide association studies (GWAS) have identified more than 25 variants near genes uniquely expressed in microglia that are predicted to increase susceptibility for AD. In light of this complexity, the mouse represents a critical model system to dissect the role of microglia and other glia in AD.

There has been large debate regarding the alignment of mouse microglia to human microglia in terms of identity, diversity and function. With the more widespread use of single-cell sequencing technology, a number of groups have suggested that the species difference is too great for conclusions drawn from mouse models to inform our understanding of human microglia^1,2^. Central to this argument is the discovery and description of a specific class of microglia in the mouse, disease-associated microglia (DAM)^3^. Based upon the current data it is unclear whether the presence or absence of DAM in human AD patients is the result of differences in tissue collection, extraction of cells, genetic diversity of patients, sub-type of AD presented in donors or even the relevant disease^1,4^. Recent work has demonstrated that single-nucleus RNA sequencing of stored human tissue fails to detect differences in microglia activation between AD and controls^5^, further complicating direct comparisons between humans and mouse models.

The vast majority of mouse microglia gene databases have been generated using the classic inbred laboratory strain, C57BL/6 (B6). Genetic complexity in the human population is expected to influence differences in, and even presence of, microglia subtypes. However, inclusion of similar genetic diversity in mouse strains has not been explored. We have taken advantage of genetically diverse wild-derived mouse strains that exhibit natural genetic variation in AD risk genes^6^. As these wild-derived strains CAST/EiJ (CAST), WSB/EiJ (WSB) and PWK/PhJ (PWK) were captured from the wild from different geographical regions for laboratory use, their genomes are closer to recapitulating the diversity of genetic variants that would exist in the natural world. We have already demonstrated that these wild-derived strains show variation in their baseline number of myeloid cells, and so it is plausible that differences in both adaptive and innate immunity may confer resilience or susceptibility to neurodegenerative diseases. The inclusion of human mutations associated with amyloid pathology, *APP*_*swe*_ and *PS1*_*de9*_ (*APP/PS1*) further highlighted strain-specific differences in neuroinflammatory responsiveness. For example, CAST.*APP/PS1* demonstrated a hyperproliferative phenotype with the highest density of microglia around plaques and, WSB.*APP/PS1* showed the fewest plaque-associated microglia. Brain transcriptional profiling across the four strains revealed that in comparison to B6, PWK mice exhibited a robust increase in microglial gene expression whereas WSB showed a greatly reduced response despite significant pathology. This strong correlation between microglia phenotypes and neuronal cell loss suggests the wild-derived AD panel provides a unique opportunity to understand the role of microglia biology on neurodegeneration. To achieve this, we developed a single myeloid cell data resource from wild-derived mouse strains that supports the importance of including natural genetic variation to study microglial biology in AD.

## Results

### Natural genetic variation shapes the transcriptome landscape of myeloid cells

In order to understand microglia diversity present in wild-derived strains compared to B6, we performed single-cell RNA sequencing (scRNA-seq) on brain myeloid cells isolated from female 9-month B6.*APP/PS1*, CAST.*APP/PS1*, PWK.*APP/PS1* and WSB.*APP/PS1* and wild-type (WT) controls. We focused on female mice as they showed the most variation in AD-relevant phenotypes at this age compared to males. Briefly, animals were perfused, and mechanical dissociation^7^ was performed on brains to obtain a single-cell suspension for myeloid cells enrichment using magnetic-activated cell sorting with CD11b-microbeads. All steps were performed on ice or at 4°C to minimize tissue dissociation-related microglia activation^8^. Single myeloid cell RNA libraries were generated using 10x Genomics v3 chemistry and sequenced by Illumina Nova-seq S2 sequencer (see Methods). Fastq files were aligned to customized strain-specific genomes by scBASE^9^ and gene counts were estimated by emaze-zero^10^(**Fig. 1a)**. Gene count matrix and downstream clustering analysis was processed using the Seurat package after removing low-quality cells and non-myeloid cells contamination (**Extended Data Fig. 1a-b**). The resulting UMAP plot showed brain myeloid cells clustered primarily according to their strain background (**Fig. 1b**), indicating strain drives the largest variation in myeloid cell gene expression. To determine and compare myeloid cell types, gene expression profiles were integrated using canonical correlation analysis (CCA)^11^. After integration, myeloid cells from each strain were clustered together, allowing for direct comparison (**Fig. 1c-d**). A total of four major myeloid cell clusters were defined, microglia (96.2%), perivascular macrophages (1.5%), monocytes (1.5%) and neutrophils (0.8%) based upon commonly used marker genes including *Itgam, Tmem119, P2ry12, C1qa, Ptprc, Mrc1, Cd74, Itgal, S100a4*, and *S100a9*^3,12^ (**Fig. 2**).

**Fig. 1.**
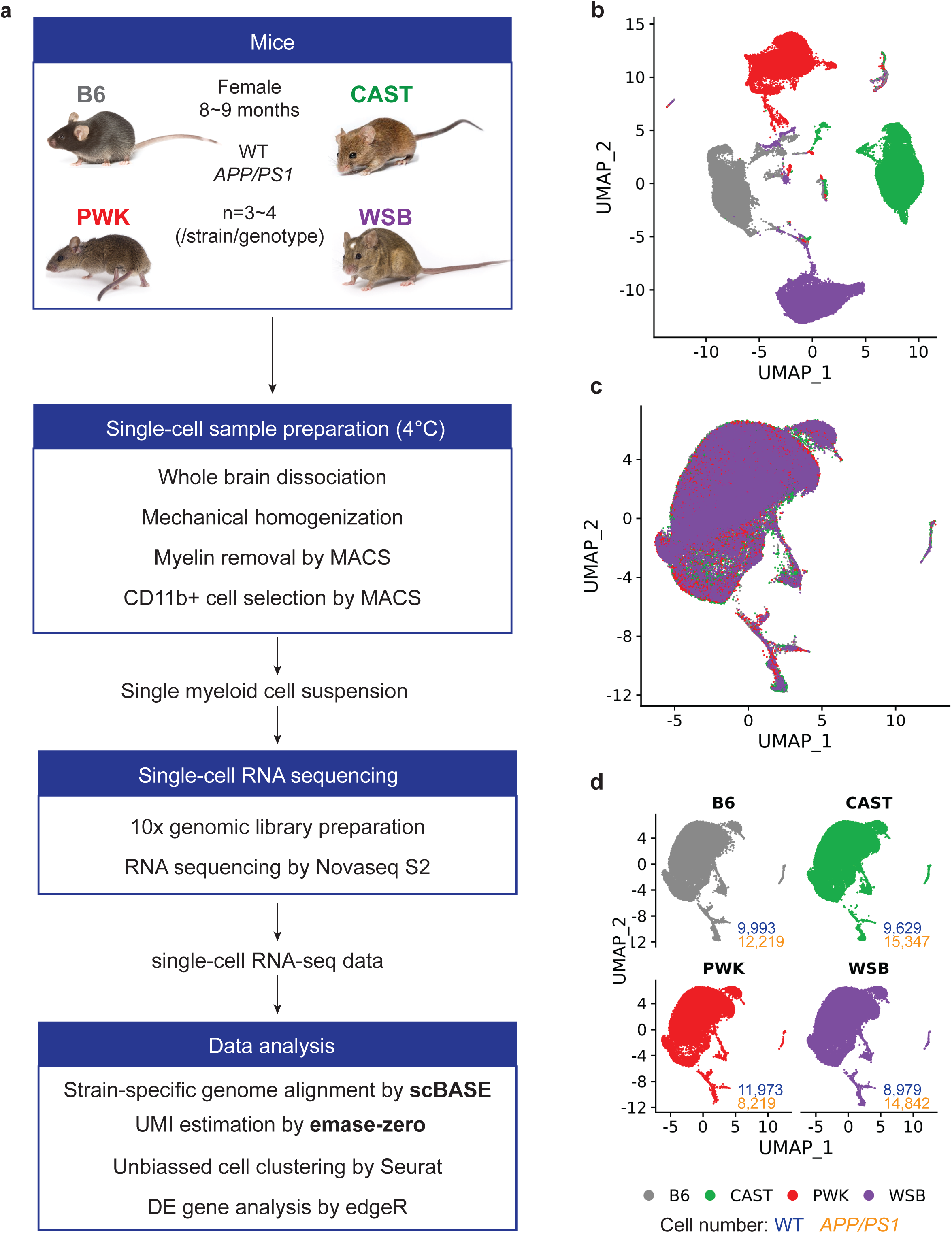
A single-cell atlas revealed natural genetic variation shapes the transcriptome of brain myeloid cells. **a**, Overview of the experimental strategy. **b**, Raw cell map of brain myeloid cells in WT and *APP/PS1* of B6, CAST, PWK and WSB mice. UMAP of 92,698 single brain myeloid cell profiles in all groups of mice (n = 3-4/genotype/strain, 29 mice in total), without data integration on strain. **c**,**d**, Strain-integrated cell map of brain myeloid cells of WT and AD of B6, CAST, PWK and WSB mice. UMAP of 91,201 single brain myeloid cell profiles in all groups of mice with merged view (c) and split views based on strain (d), with canonical correlation analysis (CCA)-based integration on strains.

**Fig. 2.**
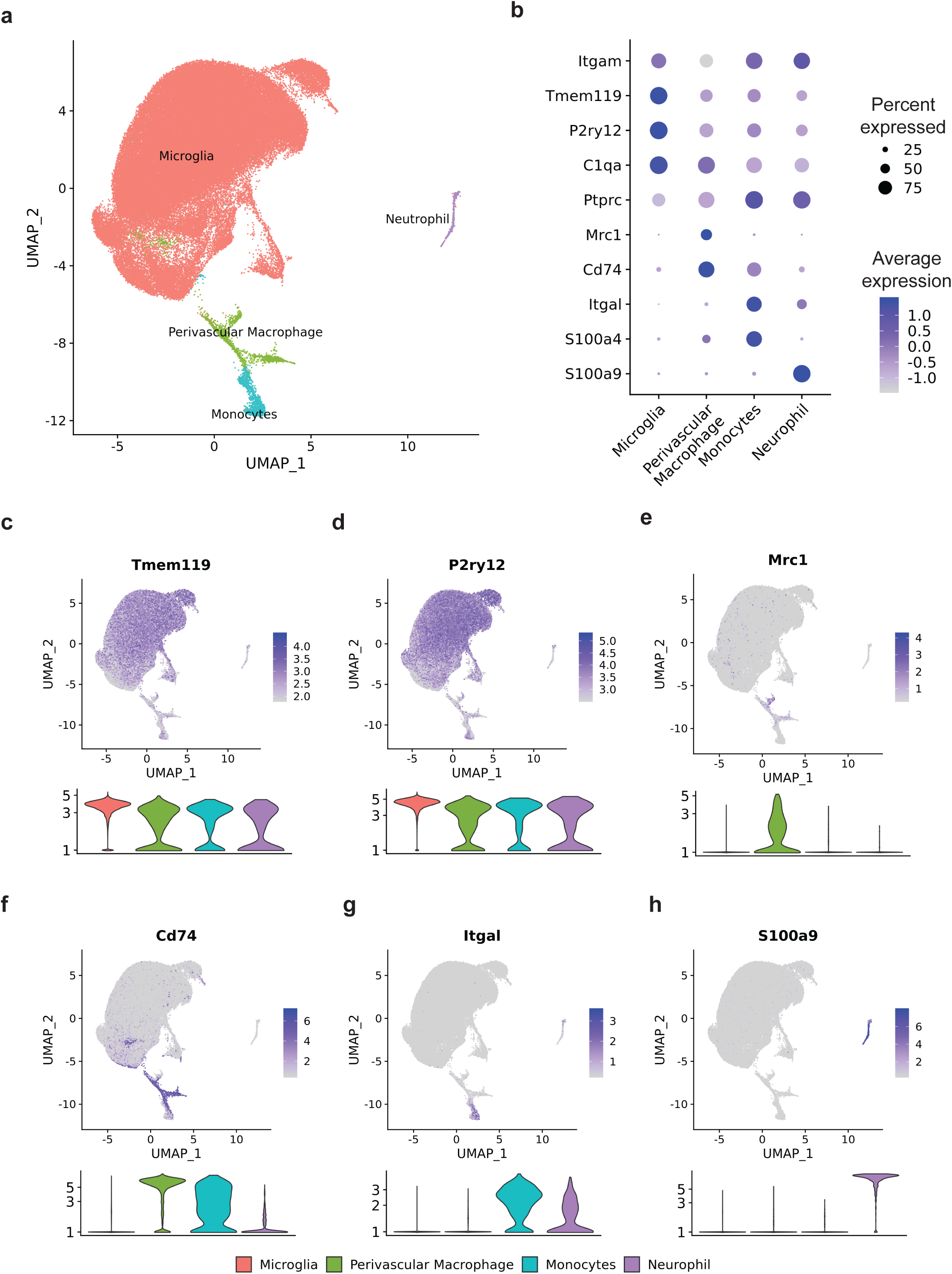
Clustering and annotation of brain myeloid cell types in B6 and wild-derived mice. **a**, Single-cell profiling revealed 4 major distinct myeloid cells including microglia, perivascular macrophages, monocytes and neutrophils. The integrated UMAP showing 91,201 single brain myeloid cell profiles in all groups (29 mice, same as Fig. 1c) colored by myeloid cell types. **b**, Dot plot showing the classical marker genes for myeloid cell types with their percent expressed and average expression. **c-h**, UMAP and violin plots featuring myeloid cell marker genes *Tmem119, P2ry12, Mrc1, Cd74, Itgal*, and *S100a9*.

### Defining microglia subtypes in genetically diverse mouse strains

Given microglia were the most common myeloid cell identified, a second round of clustering was performed to more accurately define microglia subtypes. Thirteen microglia clusters were annotated based on relative expression levels of marker genes such as *Tmem119, Cx3cr1, Cst7, Clec7a, Apoe, Ifitm3, Hexb, C3ar1* and *Stmn1* (**Fig. 3a-b, Supplementary Table 1**). Cluster number was assigned and ordered by the overall abundance of cells that exhibit similar gene expression. Clusters 0-5 were the most abundant, appeared consistent with ‘homeostatic’-like microglia, and were pooled (**Extended Data Fig. 3a-b**, collectively referred to as cluster H or homeostatic microglia). One additional cluster (Cluster 8) showed similarities to cluster H, but many of the homeostatic marker genes were expressed at a significantly higher level including *Tmem119, Hexb, Cd81* and *Cst3* (**Fig. 3b,e**) and were termed hyper-homeostatic. Clusters 6 and 12 were identified as disease-associated microglia (DAM) based on *Cst7, Lpl, Clec7a* (high expression) and *Cx3cr1* (low expression) (**Fig. 3b-c**)^3^. The two DAM clusters differed primarily in expression of homeostatic marker genes such as *Cx3Cr1, Csf1r, Tgfbr1 and Tgfbr2* as well as higher expression of *Tyrobp, Cst7* in cluster 12 (**Fig. 3a-b, Extended Data Fig. 4a**). Furthermore, while both exhibited a high ribosomal gene signature suggestive of enhanced translational activity compared to cluster H, this was greatest in cluster 12 (p. adj < 10^−16^, **Extended Data Fig. 5, Supplementary Table 2).** The high ribosomal gene feature was also found in cluster 9 which was in close proximity to cluster 12 DAM (**Fig. 3a-b, Extended Data Fig. 5)**. The predicted function of this cluster remains to be discovered. Cluster 7 was identified as interferon-responding microglia (IRM), defined by specific expression of *Ifit3, Ifitm3* and *Irf7* (**Figure 3b, d)**. Cluster 10, a relatively small cluster, expressed high levels of *Ccl3, Ccl4* and *C3ar1* (**Fig. 3b, Extended Data Fig. 6a**). Recent work identified a similar small population of microglia present during development that expand with aging, or in the context of injury^13^, amyloidosis and tauopathies^14^. Cluster 11 was identified as proliferative and was enriched for *Stmn1* (**Fig. 3b, Extended Data Fig. 6b**). No clusters showed dramatic enrichment of immediate early genes as an indication of ex vivo microglia activation^8^ (**Extended Data Fig. 2**). Importantly, although some of the eight clusters were ascribed a putative function (e.g. hyper-homeostatic, DAM, IRM, proliferative), the precise roles of all clusters in AD and other neuroinflammatory contexts remain to be discovered.

**Fig. 3.**
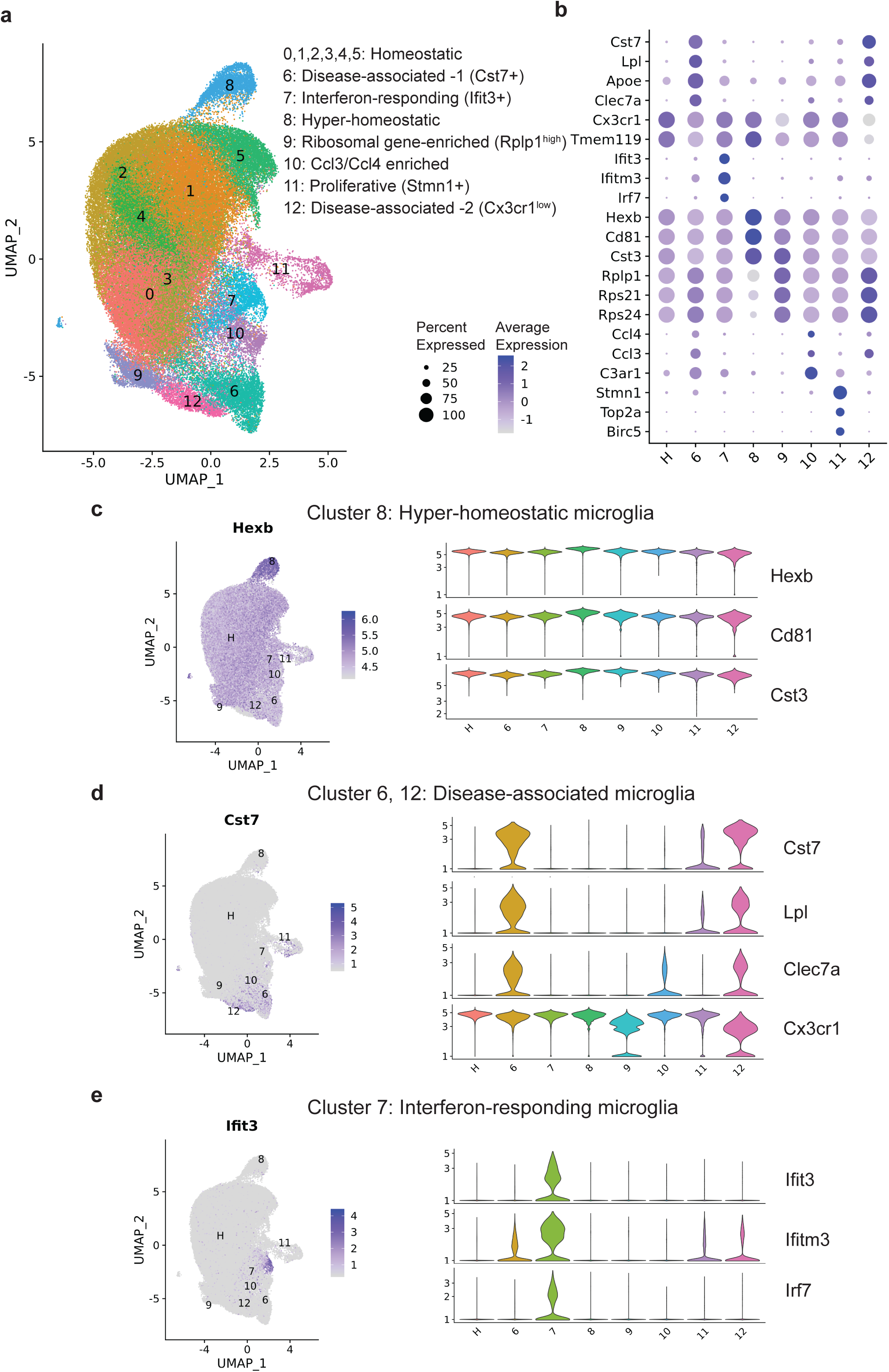
Clustering and annotation of microglia subtypes in B6 and wild-derived mice. **a**, UMAP plot showed 87,746 strain-integrated microglia from all 29 mice (20,732 from B6, 24,124 from CAST, 19,702 from PWK and 23,188 from WSB), reflecting diverse microglia subtypes including homeostatic (cluster 0-5), disease-associated (cluster 6, 12), interferon-responding (cluster 7), hyper-homeostatic (cluster 8), ribosomal gene-enriched (cluster 9), Ccl3/Ccl4-enriched (cluster 10), proliferative microglia (cluster 11). **b**, Dot plot showing the classical marker genes for myeloid cell types with their percent expressed and average expression. **c-e**, UMAP and violin plots highlighting microglia subtype marker genes for hyper-homeostatic (c), disease-associated (d) and interferon-responding (e) microglia.

### Wild-derived strains reveal transcriptomic variation in microglia subtypes

Variation in microglia subtypes was identified by comparing percent of cells in each cluster between each strain/genotype (**Fig. 4a, Supplementary Table 3, Extended Figure 3c-d**) as well as using trajectory inference analysis where all eight subtypes were plotted across pseudotime (**Fig. 4b**). The percent of homeostatic microglia (cluster H) was significantly decreased in *APP/PS1* mice of B6, CAST and PWK strains compared to their WT counterparts. However, this was not the case for WSB.*APP/PS1* mice, which showed a similar abundance of homeostatic microglia to WSB WT (**Fig. 4c**).

**Fig. 4.**
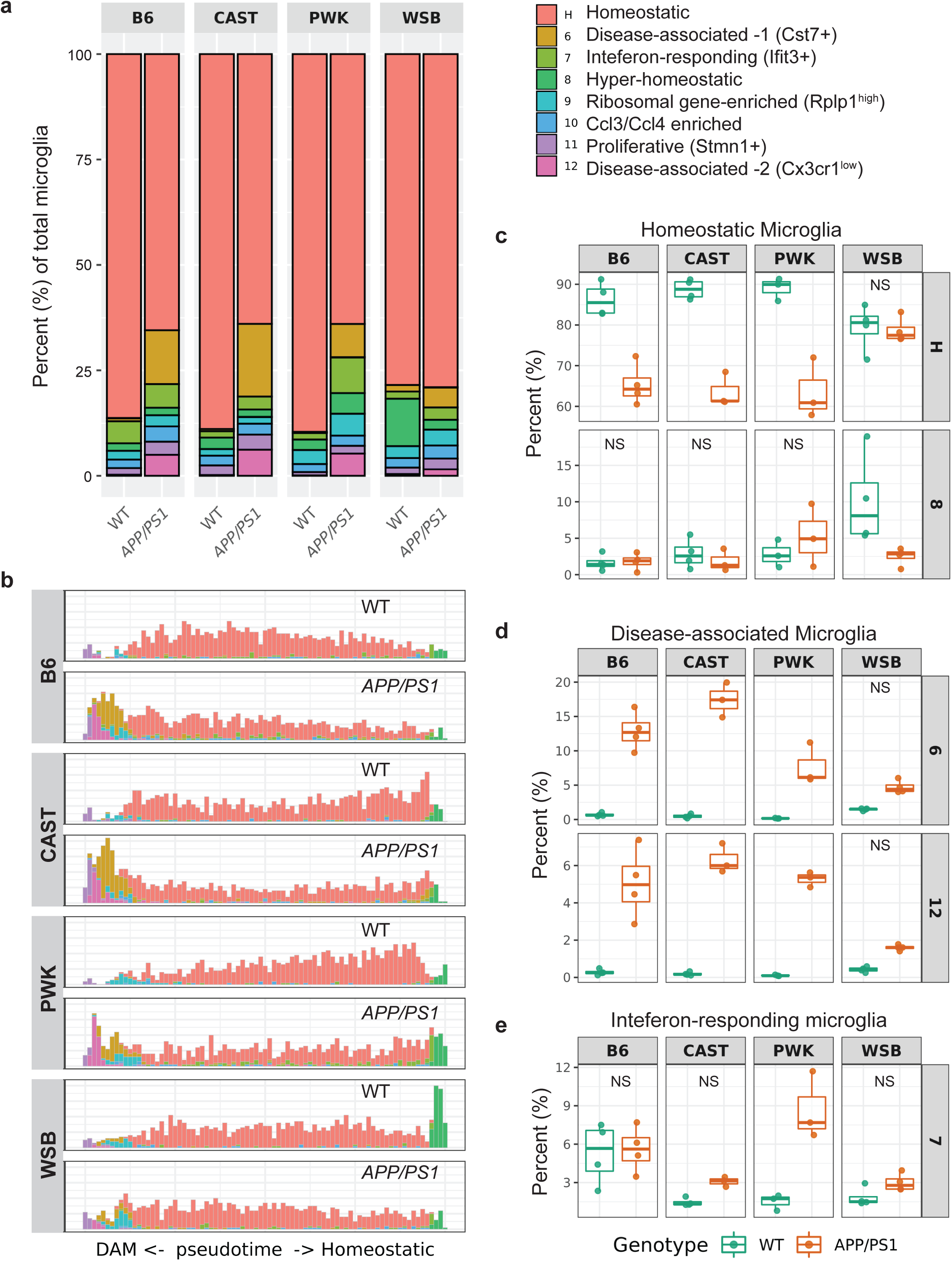
Abundance of microglia subtypes is highly variable in B6 and wild-derived strains. **a**, Abundance percentage of microglia subtypes in WT and *APP/PS1* mice of B6, CAST, PWK and WSB. **b**, Histogram of the pseudotime (the first dimension of the diffusion map) showing the distribution of 1000 microglia randomly sampled from each group. **c-e**, Box plots showing the percent of homeostatic (cluster H) and hyper-homeostatic (cluster 8) microglia (c), disease-associated microglia (d) and interferon-responding microglia (e) in all group of mice. Strain, genotype and strain by genotype effects were assessed by 2-way ANOVA followed by Tukey’s post hoc test. All comparisons (comparing WT and *APP/PS1* within each strain for a given cluster) were significant (adjusted p value, p.adj < 0.05) except for those labelled with NS (not significant, p.adj ≥ 0.05). Detailed p.adj values and confidence intervals for within and cross strain/genotype comparisons were reported in Supplementary Table 3.

Trajectory inference analysis predicted a transition of homeostatic microglia to hyper-homeostatic microglia and DAM (**Fig. 4b**). This suggests that differences in homeostatic clusters between strains may correspond to differences in transitions to other subtypes. There was a significantly greater percentage of hyper-homeostatic microglia (cluster 8) in WSB WT mice compared to other WT strains (**Fig. 4c)**, and these were largely absent in WSB.*APP/PS1*. Importantly, while the percent of DAM (clusters 6 and 12) was robustly increased in *APP/PS1* mice of B6, CAST and PWK compared to their WT counterparts, there was no significant increase in WSB.*APP/PS1* mice compared to their WT control (**Fig. 4d).** In addition, the percent of IRM (cluster 7) differed between strains. PWK.*APP/PS1* mice exhibited a significantly greater proportion of IRM in comparison with PWK WT mice. This significant *APP/PS1*-dependent increase was not observed in other strains (**Fig. 4e**). B6.*APP/PS1* was the only strain to show a genotype-specific increase in the percentage of *Ccl3/Ccl4*-enriched cells (cluster 10, **Extended Data Fig. 6d**). Finally, B6.*APP/PS1* and CAST.*APP/PS1* showed a significant increase in the percent of proliferative microglia (cluster 11) compared to their WT counterparts (**Extended Data Fig. 6e**). Collectively, these analyses show genetic diversity resulted in abundance differences in microglial subtypes in our wild-derived AD panel compared to B6.

### Transcriptome diversity in multiple genetic background reveals the functional diversity of microglia subtypes

Despite consistency of expression of marker genes within microglial clusters across strains, initial clustering suggested strain-specific gene expression differences (**Fig. 1b**). These differences could be critical for the variation we observed in amyloid-induced outcomes^6^. Given their previous association to aging and AD, we chose to focus on homeostatic (cluster H), DAM (cluster 6), IRM (cluster 7) and *Ccl3*/*Ccl4-*enriched (cluster 10) subtypes. Given the greater number of cells in cluster 6 compared to 12 (**Fig. 4d**), we focused comparisons on DAM cluster 6. The homeostatic clusters from wild-derived strains were compared to the B6 homeostatic cluster, and unique and overlapping differentially expressed (DE) genes determined. These data indicate substantial gene expression differences in homeostatic microglia across the strains (**Fig. 5a-e, Supplementary Table 4**). Next, we used Diseases and Functions analysis in Ingenuity Pathway Analysis (IPA) to predict how strain-specific differences in gene expression leads to differences in microglia function (**Fig. 5f**), followed by Regulatory Effect (RE) analysis to predict the upstream regulator(s) that may drive such functional differences for each strain (**Fig. 5g-h**). As an example, diseases and functions predicted to be downregulated in PWK compared to other strains related to ion channels (‘Flux of divalent cations’, ‘Flux of ion’, ‘Ion homeostasis of cells’, ‘Flux of inorganic cation’ and ‘Flux of Ca2+’, **Fig. 5g**). This included downregulation of *Clec7a, Cybb, Wnt4* and *Ctsb* mediated by predicted upstream regulators L2HGDH, PRKCA, Saa3, Klra7 and TNNI3. Homeostatic microglia are considered to be in a sensing state^15^, equipped to detect environmental changes in order to respond to a variety of stimuli. At the center of this transformation is identification of several surface channels and receptors that are critical for entry of calcium ions^16,17^. Thus, PWK are predicted to be a novel strain in which to understand differences related to this process. As a second example, diseases and functions predicted to be downregulated in homeostatic microglia in WSB compared to the other strains centered on myeloid cell number (‘Quantity of cells’, ‘Stimulation of cells’, **Fig. 5f**). This includes downregulation of *Ccr2, Il1b, Tnf* and *Il6* mediated by the upstream regulator LGALS3. This supports previous work that shows WSB have ∼50% fewer microglia than B6^6^.

**Fig. 5.**
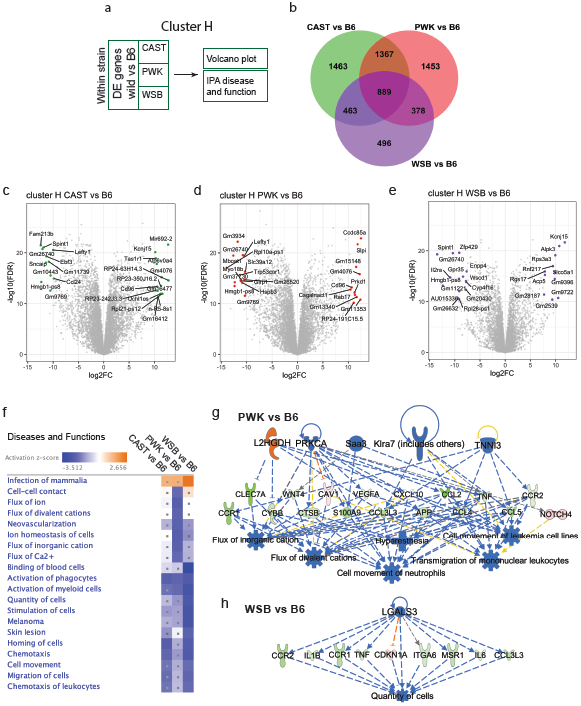
Strain-specific gene expression in homeostatic microglia. **a**, Schema for strain-specific DE gene analyses for homeostatic microglia. DE genes comparing wild-derived strains to B6 (FC > 2, FDR < 0.05) were analyzed through a venn diagram, volcano plot, and IPA. **b**, Venn diagram displaying the numbers of overlapping DE genes resulting from multiple comparisons of CAST vs B6, PWK vs B6 and WSB vs B6. **c-e**, Volcano plots showing DE genes comparing CAST vs B6 (c), PWK vs B6 (d) and WSB vs B6 in homeostatic microglia (e). Top 25 DE genes based on fold change (FC) and false discovery rate (FDR) were colored and labelled. **f**, Heat map summarizing top 20 significantly enriched terms of Diseases and Functions based on DE genes from comparisons of wild-derived vs B6 mice [corrected p-value using Benjamini-Hochberg false discovery rate (pval-BH) < 0.05, and |z-score| ≥ 2]. **g-h**, Examples of Regulatory Effects for PWK vs B6 (g) and WSB vs B6 (h) highlighting upstream regulators (top), downstream targets (middle) and diseases and functions (bottom). The orange and blue colors indicate the predicted up- or down-regulation of an upstream regulator or a Disease and Function term for a given comparison of wild-derived strain to B6. The red and green colors show the up- or down-regulation of the downstream targets as DE genes in a given comparison of wild-derived strain to B6.

DAM and IRM are the more prominent microglia subtypes previously implicated in AD ^3,13,18^. Therefore, we sought to (i) identify differences in cluster-defining genes between strains and (ii) identify wild-derived cluster differences in comparison with B6 (**Fig. 6-7**). DAM-specific DE genes were identified by comparing cluster 6 to cluster H within each strain (**Fig. 6b, Supplementary Table 5)**. While some genes were consistently DE across all strains (such as the DAM marker genes *Cst7, Lpl, Clec7a* and *Apoe*), many were DE in three, two or even only one strain. For instance, CAST and PWK had 126 and 50 unique DE genes respectively. To better understand the degree of variation in DAM cluster-defining genes, a Spearman Correlation was performed based on the fold change of DE genes (**Fig. 6c**) and differences were visualized (**Fig. 6d-f**). This indicated that CAST DAM were the least similar to B6 (correlation coefficient = 0.76), followed by PWK (0.79) with WSB being the most similar (0.89). Comparison of wild-derived DAM to B6 DAM identified DE genes (**Figure 6g-i, Supplementary Table 4**) that were predicted to impact a number of diseases and functions (**Figure 6j**). For example, CAST showed a significant activation of genes related to ‘Cellular Infiltration of Mononuclear Leukocytes’, regulated by IL3 (**Figure 6k**). IL3 is a growth factor and cytokine involved in homing microglia to plaques and is thought to be neuroprotective^19^. IL3 enrichment is driven by the upregulation of genes including *Vcam1, Cd14* and *Casp3* in CAST DAM compared to B6 DAM. A second example of differences between strains is the network centered around ‘Binding of Endothelial Cells’. Genes shown to be necessary for binding of myeloid cells to endothelial cells, such as integrins (e.g. *Itga6* and *Itgal*), are significantly downregulated in WSB DAM compared to B6 DAM (**Figure 6l**). Vascular dysfunction has more recently been identified as a risk factor for many cases of AD and related dementias. Interestingly, WSB.*APP/PS1* were the only strain in which there was no increase in DAM, and show the highest levels of cerebral amyloid angiopathy (CAA), vascular leakage and neuronal loss^6^. These data suggest that WSB.*APP/PS1* may be an important model to understand the interplay between immune function and vascular changes in relation to amyloid pathology and neurodegeneration.

**Fig. 6.**
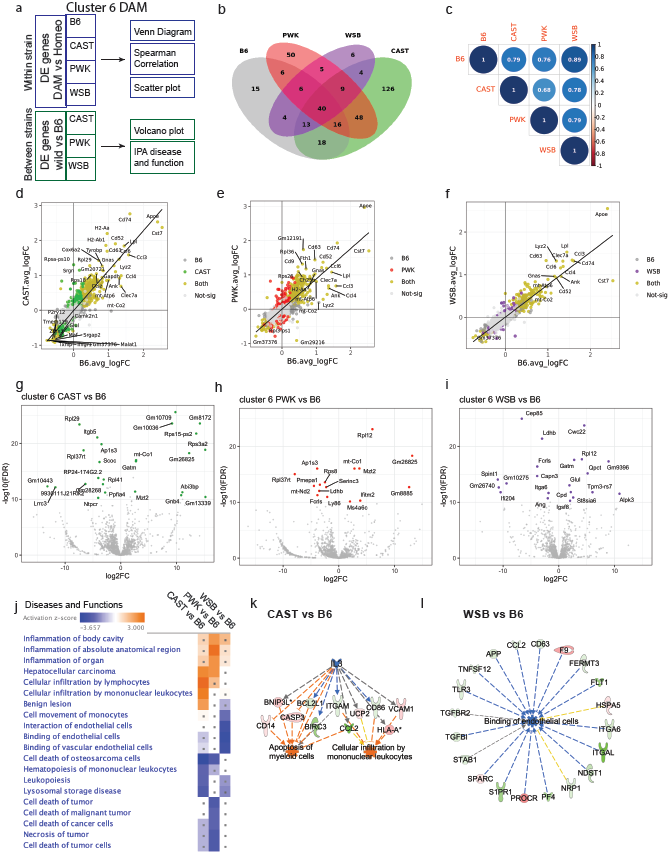
Strain-specific gene expression in disease-associated microglia (DAM). **a**, Schema for strain-specific DE gene analyses in DAM (cluster 6). DAM marker genes comparing DAM to homeostatic microglia (FDR < 0.05) in each strain were analyzed through a venn diagram, spearman correlation and scatter plots. DE genes comparing wild-derived IRM to B6 IRM were analyzed in a volcano plot and IPA. **b**, Venn diagram showing the overlap of the numbers of DAM marker genes across strains. **c**. Spearman correlation matrix based on fold change (FC) of top DAM marker genes (|log2FC| > 0.5) between strains showing the correlation coefficients. All correlations are significant (p < 0.05). **d-f**, Scatter plots showing DAM marker genes between CAST vs B6 (d), PWK vs B6 (e) and WSB vs B6 (f). B6 (dark grey): significant only in B6 but not in wild-derived strains; CAST (green), PWK (red), WSB (purple): significant only in CAST, PWK or WSB but not in B6; Both (yellow): significant only in both wild-derived and B6 mice. **g-i**, Volcano plots showing DE genes comparing CAST vs B6 (g), PWK vs B6 (**h**) and WSB vs B6 (i). Top 25 gene DE genes based on FC and FDR were labelled. **j**, Heat map summarizing top 20 significantly enriched Diseases and Functions (IPA) terms based on DE genes from comparisons of wild-derived vs B6 mice (pval-BH < 0.05, |z-score| ≥ 2). **k**, Example of a Regulatory Effect for CAST vs B6 highlighting network of ‘Apoptosis of myeloid cells’ and ‘Cellular infiltration by mononuclear leukocytes’. **l**, Example of a Regulatory Effect for WSB vs CAST highlighting the of ‘Binding of endothelial cells’. The color code is as described in Fig.5 g-h.

We performed the same series of analyses on IRM as described for DAM (**Fig. 7a**). Comparison of DE gene expression in IRM versus homeostatic microglia highlighted a core set of 24 genes that were DE in all strains and included top marker genes *Ifitm3, Ifit3* and *Irf7* (**Fig. 7b, Supplementary Table 5**). There were also DE genes that were unique to specific strains such as CAST (e.g. 8 DE genes comparing CAST IRM to CAST homeostatic microglia). Although there was a significant correlation in all comparisons (p < 0.01), the correlation strength between strains was more variable in IRM (**Fig. 7c-f**) than in DAM (**Fig. 6c-f**). DE genes were identified comparing wild-derived IRM and B6 IRM (**Fig. 7g-i, Supplementary Table 5**) that were predicted to differentially affect multiple diseases and functions (**Figure 7j-k**). For instance, activation of a network that relates to ‘Liver Damage’ and ‘Immune response of cells’ were unique to CAST. Interestingly, genes that comprise these networks, *Irf7, Birc3, Tnfsf10, Il6, Serpine1* and *Tab1* (predicted to be the upstream regulator), have been shown to be differentially expressed in brains of AD patients^20^, and identified as targets for therapeutics^21-27^. Based on our data, CAST, but not B6, would be ideal to assess drug targets that engaged genes in this network. Other networks identified as down-regulated in CAST compared to other strains was ‘Activation of lymphocytes’ and ‘Antimicrobial response’. Interferons are a group of cytokines secreted in response to stress or viral infection and are associated with autoimmune diseases. Patients with HIV-induced dementia exhibit increases in interferon activation^28^, and the viral theory of AD has recently made a resurgence^29^. Upstream regulators *NLRX1, NKX2-2* and *TLR8* are all related to Type 1 interferon-triggering components such as *STAT1* and *MYD88*. Nucleic acid containing amyloid fibrils can potently induce this cascade^18^. Furthermore, increases in NLRX1, cytoplasmic NOD-like receptors localized to the outer membrane of mitochondria, have been associated with increased production of reactive oxygen species^30^. This suggests that strategies that compare CAST.*APP/PS1* (low expressers) with PWK.*APP/PS1* (high expressers) would be appropriate to parcel this relationship between viral immune pathways and AD.

**Fig. 7.**
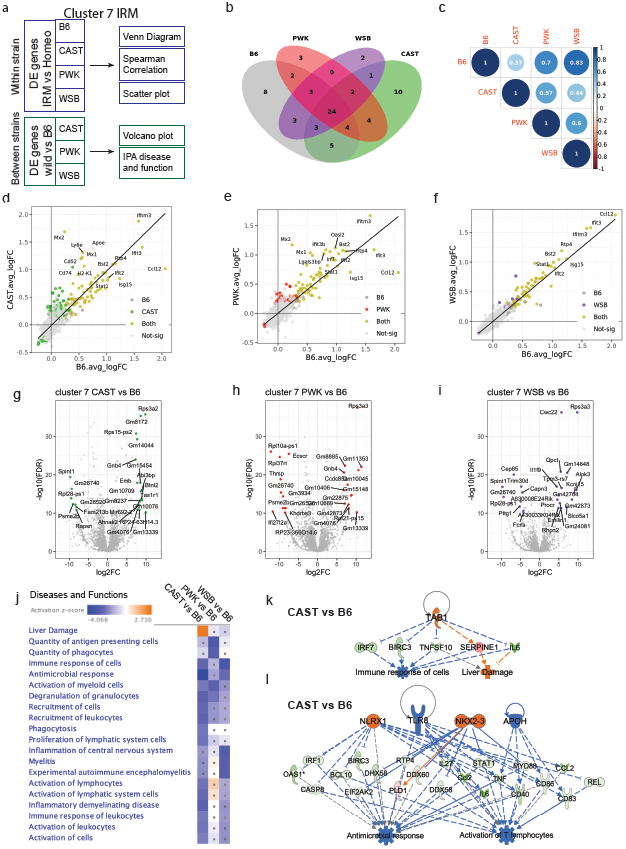
Strain-specific gene expression in Interferon-responding microglia (IRM). **a**, Schema for strain-specific DE gene analyses in IRM (cluster 7). IRM marker genes comparing IRM to homeostatic microglia (FDR < 0.05) of each strain were analyzed through a venn diagram, spearman correlation and scatter plots. DE genes comparing wild-derived IRM to B6 IRM were subjected to volcano plot and IPA. **b**, Venn diagram showing the overlap of the numbers of IRM marker genes across strains. **c**. Spearman correlation matrix based on FC of top IRM marker genes (|log2FC| > 0.5) between strains showing the correlation coefficients. All correlations are significant (p < 0.05). **d-f**, Scatter plots showing IRM marker genes between CAST vs B6 (d), PWK vs B6 (e) and WSB vs B6 (f). B6 (dark grey): significant only in B6 but not in wild-derived strains; CAST (green), PWK (red), WSB (purple): significant only in CAST, PWK or WSB but not in B6; Both (yellow): significant only in both wild-derived and B6 mice. **g-i**, Volcano plots showing DE genes comparing CAST vs B6 (g), PWK vs B6 (h) and WSB vs B6 (i). Top 25 gene DE genes based on FC and FDR were labelled. **j**, Heat map summarizing top 20 significantly enriched terms of Diseases and Functions based on DE genes from comparisons of wild-derived vs B6 mice (pval-BH < 0.05, |z-score| ≥ 2). **k**, Example of a Regulatory Effects for CAST vs B6 highlighting network of ‘Immune response of cells’ and ‘Liver damage’. **l**, Example of a second Regulatory Effect for CAST vs B6 highlighting ‘Antimicrobial response’ and ‘Activation of T lymphocytes’. The color code is the same as described in Fig.5 g-h.

Finally, comparison of the *Ccl3*/*Ccl4-*enriched clusters between wild-derived strains and B6 identified differences related to interactions with other immune cells **(Extended Data Fig. 7)**. *Ccl3*/*Ccl4-*enriched microglia have been previously localized to the center of active demyelinating lesions in multiple sclerosis patients^13^, and are suggested to signal to peripheral immune cells. Interestingly, in comparison with B6, CAST show a downregulation in pathways relevant to ‘Multiple sclerosis’, ‘Inflammatory demyelinating disease’ and ‘Extravasation of cells’ **(Extended Data Fig. 7j-k)**. This indicates that neurodegenerative processes in CAST may be independent of damage caused by infiltrating immune cells. Collectively, these analyses confirm significant variation in genes in disease-relevant pathways between wild-derived and B6 in multiple microglial subtypes.

### Human GWAS AD genes are differentially expressed in microglia subtypes

Variation in microglia-relevant genes are differentially associated with AD risk. However, previous studies to determine roles of GWAS genes in AD have primarily been limited to the B6 genetic background. Therefore, we aimed to determine whether our wild-derived AD panel provided an enhanced platform to study human-relevant AD genes using a panel of 54 GWAS genes identified in two recent meta-analyses (**Supplementary Table 6**) ^31,32^. A total of 36 microglia-relevant genes were detectable across our panel. Nineteen of the 36 genes (52%) were DE (FDR < 0.05) in at least one cluster comparing wild-derived strains to B6 (**Fig. 8a**). Genes could be DE in only one cluster of one strain (e.g. *Adam10* in cluster 8, CAST vs B6; *Bin1* in cluster 8, PWK vs B6; *Inpp5d* in cluster 6, PWK vs B6; and *Pilra* in cluster 6, WSB vs B6), while other genes were DE in multiple clusters within a specific strain (e.g. *Ptk2b* and *Ndufa1* in CAST; *App* and *Sorl1* in PWK). *Scimp* and *Apoe* were DE in at least one cluster in all wild-derived strains compared with B6. The expression in WT and *APP/PS1* mice across the four strains was then determined for cluster H, DAM and IRM (**Figure 8b**). This further highlighted strain and genotype specific differences in GWAS genes. For instance, *Sorl1* was expressed in many more cells in IRM (cluster 7) from PWK mice compared to B6, CAST and WSB. Moreover, the relative expression level of *Sorl1* was increased significantly in PWK.*APP/PS1* compared to PWK mice. Therefore, these data further support the use of specific or contrasting wild-derived strains for more extensive and informative functional studies of AD GWAS genes.

**Fig. 8.**
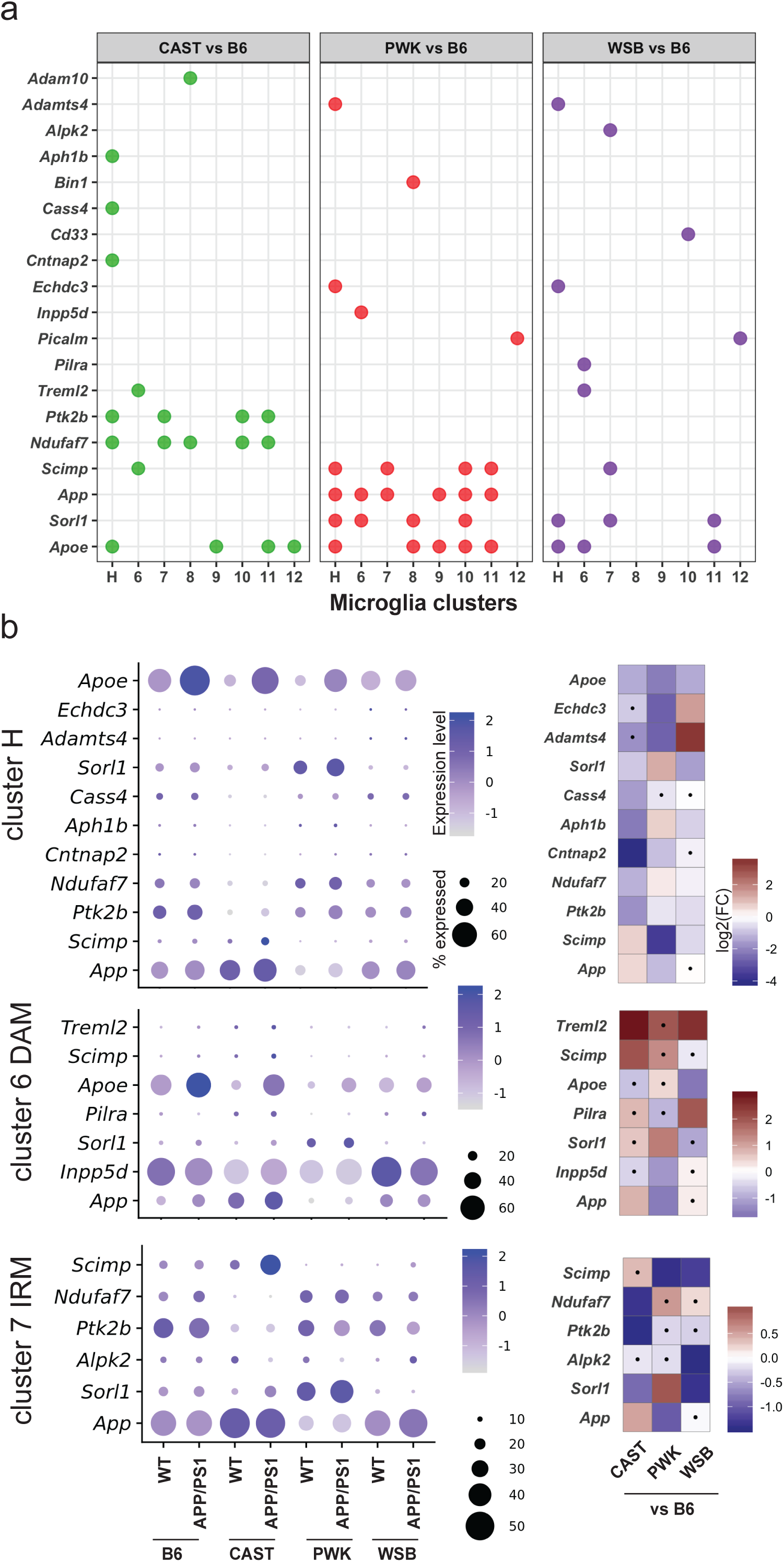
Microglia subtypes from wild-derived strains show differential expression of AD GWAS genes. **a.** Nineteen AD GWAS genes were DE comparing wild-derived strains to B6 for all eight microglia clusters. Dot indicates the gene was significantly DE (FDR < 0.05) comparing CAST, PWK, or WSB to B6 in a given cluster. **b.** Dot plot (left) showing the percent of cells expressed and the expression levels of significant strain-specific DE genes in homeostatic (cluster H), DAM (cluster 6) and IRM (cluster 7) across all groups. Heatmap (right) highlighting fold change (log2FC) of the corresponding gene expression comparing CAST, PWK and WSB to B6 in cluster H, 6 and 7. Dots in the heatmap indicate the fold change for a given comparison is not significant (FDR ≥ 0.05).

## Discussion

Single-cell sequencing of microglia from wild-derived and B6 mouse strains revealed that natural genetic variation led to significant differences is gene expression profiles that would significantly impact microglia biology, leading to an inherently different neuroimmune environment in healthy and diseased states. It is likely that these observed variations in microglia states between strains influences, or is influenced by, other cell types including astrocytes, endothelial cells and neurons. These data begin to reveal the full potential of using mouse genetic diversity to unravel the complexity of neuroinflammation in AD. In this study, microglia from female mice at one age (8 months) were profiled. In addition to all the strain-specific and strain-by-genotype specific changes observed in this dataset, sex-, brain region- and/or age-specific changes are still to be determined. These differences will be important as microglia, and more generally neuroinflammation, are central to brain health throughout aging, and in the context of comorbidities for AD such as obesity, cardiovascular disease, diabetes, and viral infection.

Differences across wild-derived strains in microglia subtypes often showed downregulation of specific biological pathways in comparison to B6. While B6 has been used across biomedical research for practical and historical reasons, such work may be inherently biased to neuroimmune responses driven by a singular genetic context with limited translation to humans. For example, B6 (as well as other commonly used strains such as DBA/2) carries a mutation in the *P2rx7* locus that severely impairs important functions of this receptor. This is thought to influence critical steps relating to induction of apoptosis and cytokine secretion. In contrast, wild-derived strains carry the ‘natural’ variant also present in humans^33^. Another key consideration is that previous microglia sequencing projects have used the 5XFAD model. There are two versions of this model, one congenic on B6 (JR# 34848), and the other more commonly used B6.SJL mixed genetic background (JR# 34840). SJL mice carry the *Trem2*^*S148E*^ mutation, which means that 5XFAD could be heterozygous, homozygous or wildtype for this mutation, influencing microglia function differently within the same study or across studies. Therefore, studies such as ours that incorporate genetic variation will be more relevant to the responses seen in human aging and neurodegenerative disorders.

We detected significant strain and strain-by-genotype differences. This was both in the abundance of microglia subtypes, and in gene expression that in combination with pathway analysis and neuropathology are predictive of functional differences that may be beneficial or damaging depending on stage of disease. For example, homeostatic microglia are typically defined as in a sensing state, sampling the brain environment for debris and potential pathogens. If a signal is encountered, they quickly become activated to deal with the threat. Upon resolution, microglia are expected to revert back to their surveillance role. One theory regarding the influence of microglia to disease susceptibility is that once triggered, these microglia cannot revert back, becoming chronically ‘activated’, signaling to other local immune cells, and potentially causing damage to healthy tissue^34^. Our data predicts natural genetic variation influences the baseline responsiveness, efficiency of response, and reversion back to surveillance. Initial clustering of microglia identified 6 groups of homeostatic-like microglia that were collapsed into one cluster based upon similar marker gene expression. However, initial clustering predicted subtle but distinct functional differences that remain to be resolved. The hyper-homeostatic cluster (cluster 8) showed higher expression levels of *Cst3, Cd81* and *Hexb* compared to the homeostatic cluster. Two small *Hexb*-related clusters have been previously reported^3^; however, those clusters do not fully align with our cluster as they displayed a signature of lipid metabolism and phagocytosis with increased expression of *Apoe, Lpl* and *Cst7*. This was opposite in hyper-homeostatic microglia. Interestingly, *HEXB* has been previously associated with microglia responses in AD^35^, and is responsible for production of two enzymes, beta-hexosaminidase A and B that play a critical role in lysosomal degradation of sphingolipids. Loss of *HEXB* is associated with Sandhoff disease, a rare disorder that leads to progressive neurodegeneration of the brain and spinal cord. Therefore, hyper-homeostatic microglia may represent an intermediate, transition stage between surveillance and activation. Alternatively, given that pseudotime analysis suggested that this subtype transitioned in the opposite direction to activated subtypes like DAM and IRM, they may represent a microglia ‘reserve’ pool. This is consistent with the lower numbers of genes expressed per cell, and low levels of ribosomal genes compared to homeostatic microglia (Extended Data Fig. 5, Supplementary Table 2). WSB show the largest numbers of hyper-homeostatic microglia compared with other WT strains that are essentially non-existent in WSB.*APP/PS1* supporting the use of WSB to study these novel micoroglia subtype.

Two DAM-like clusters (clusters 6, 12) were identified based on lower expression of homeostatic markers and higher expression of *Tyrobp* in cluster 12 compared to cluster 6. However, despite these differences, no functional conseqeunces were predicted. Two previous studies have reported two subtypes of DAM. In one study, two DAM subtypes were suggested to represent *Trem2*-specific transition states^3^, while the second study predicted proinflammatory and anti-inflammatory subtypes^36^. However, neither of these were replicated in our study. The lack of replication could be due to the amyloid transgenes used. Microglia activation and amyloid accumulation has been identified as early as 6 weeks in 5xFAD mice^37,38^ but is not apparent until 4-5 months in B6.*APP/PS1* mice^39,40^. DAM populations in our wild-derived and B6 AD panel were also significantly smaller than has been previously reported in another amyloid strain, B6.*APP*_*swe*_*/PS1*_*L166P*_ ^35^ which is also an aggressive amyloid strain with plaque accumulation being observed as early as 6 weeks^41^

In contrast to the other strains, WSB.*APP/PS1* did not show a significant increase in DAM compared to their wild-type counterparts. Gene expression analyses predicted a downregulation of genes related to cellular interactions with endothelial cell in WSB compared to B6. CAA and vascular dysfunction was previously identified in WSB.*APP/PS1* mice^6^ and CAA is thought to be independent of neuroinflammation in human AD patients^42^. Recent work used the CSF1R inhibitor PLX5622 to deplete microglia in 5XFAD resulting in almost completely loss of amyloid in the parenchyma, and significant CAA and vascular leakage^43^. Therefore, WSB.*APP/PS1* may be an ideal strain to unpack the interrelationship between amyloidosis, CAA and vascular dysfunction in AD. If these differences in DAM also translate to humans, it highlights that there are likely patients who show an elevated DAM response, and patients who do not. This could partially explain controversy over the presence of DAM populations in human microglia datasets.

Our study highlights the importance of broadening our interest in microglia subpopulations beyond DAM. IRM were significantly different between strains – with only PWK.*APP/PS1* showing a significant increase compared to WT. Interferon response is a complex processes that can trigger expression of thousands of interferon stimulated genes (ISGs). Commonly, the interferon response is thought to be triggered in response to a viral infection and strain differences in viral response have been identified. CAST is uniquely susceptible to infections such as Influenza H3N2 and Monkeypox virus. In the case of Influenza H3N2, despite high viral load in the lungs, CAST exhibited an abnormal response in leukocyte recruitment^44^. Even at low inoculums of Monkeypox virus, CAST show rapid spread to all internal organs. This was shown to be directly related to deficiency in gamma interferon^45^. In AD, the interferon response can be triggered by nucleic acid (NA)-containing plaques. In our, and other studies, IRM are defined by the presence of interferon regulator gene *Irf7* as well as ISGs *Ifitm3* and *Ifit3*. In one recent study, brain samples showed the presence of IFITM3 microglia in NA+ plaques^18^. Enhancing the interferon response in a B6.5xFAD exacerbated synapse loss. In contrast, our study supports a beneficial role for IFITM3+ IRM in AD – PWK.*APP/PS1* showed increased levels of IRM compared to the other strains and are resilient to neurodegeneration at 8 months^6^. In support of this, mice deficient for IFITM3 are more susceptible to viral infection^46^. Given the multitude of outcomes downstream of the interferon response, it is critical we understand the specific role of IFITM3+ cells in AD.

In conclusion, this wild-derived AD panel offers a level of genetic and phenotypic diversity that can aid determining the role of microglia in human AD. There will continue to be debate regarding the level at which the mouse immune system should be ‘humanized’ in order to better model human immune function. However, based on our data, and with improved tools and resources, such as strain-specific gene editing protocols, and reporter and cre lines, integrating the use of wild-derived strains that exhibit variation closer to the natural world is essential.

## Online Methods

### Ethics statement

All research was approved by the Institutional Animal Care and Use Committee (IACUC) at The Jackson Laboratory (approval number 12005). Authors performed their work following guidelines established by the “The Eighth Edition of the Guide for the Care and Use of Laboratory Animals” and euthanasia using methods approved by the American Veterinary Medical Association.”

### Mouse strains and cohort generation

All mice were bred and housed in a 12/12 hours light/dark cycle on aspen bedding and fed standard 6% LabDiet Chow. Experiments were performed on four mouse strains: B6.Cg-Tg(APPswe, PSEN1dE9)85Dbo/Mmjax (JAX stock #005864), CAST.*APP/PS1* (JAX Stock #25973), WSB.*APP/PS1* (JAX Stock #25970) and PWK.*APP/PS1* (JAX Stock #25971). Generation of experimental cohorts consisted of 6 female mice (*APP/PS1* carriers and littermate wild-type controls). Due to increased pup mortality in the wild-derived strains, once determined to be pregnant, female mice were removed from the mating and housed individually. During this time, they were also given BioServ Supreme Mini-treats (Chocolate #F05472 or Very Berry Flavor #F05711) in order to discourage pup cannibalism. Animals were initially group-housed during aging and then individually housed if fighting occurred.

### Brain single myeloid cell preparation

Four mice were included in each of the B6.WT, B6.*APP/PS1*, CAST.WT, WSB.WT, and WSB.*APP/PS1* groups (n=4) and three mice were included in each of the CAST.*APP/PS1*, PWK.WT and PWK.*APP/PS1* groups (n=3). With modification from the protocol of Bohlen CJ et al ^7^, brain myeloid single-cell suspension were obtained through mechanical dissociation followed by magnetic-activated cell sorting (MACS). All procedures were performed on ice or under 4°C to avoid artificial activation of microglia during the sample preparation. Mice were anesthetized using ketamine/xylazine (10 mg ketamine and 2 mg xylazine in 0.1ml sterile pure water per 10 g body weight) and perfused using ice cold homogenization buffer [Hank’s balanced salt solution (HBSS) containing 15mM HEPES and 0.5% glucose]. Brains were quickly dissected and transferred on ice. Each brain was minced using a scalpel and then homogenized using a 15mL PTFE tissue grinder 4-5 strokes in 2mL homogenization buffer containing 320KU /ml DNaseI (Worthington. Cat# DPRFS). The cell suspension was transferred to a 50 mL tube and passed through a pre-wet (with homogenization/DNAase I buffer) 70 micron cell strainer. The filtered cell suspension was then transferred into a 15 mL tube and spun down at 500 g for 5 minutes at 4°C. The supernatant was discarded and the cell pellet was resuspended in 2 mL MACS buffer [Phosphate-buffered saline (PBS) with 0.5% BSA and 2mM Ultrapure EDTA] for myelin removal procedure. 200 μL Myelin Removal Beads II (Miltenyi Biotec #130-096-733) was added to the cell suspension and mixed gently by pipetting. The cell suspension was then divided into two 2 mL microcentrifuge tubes (1 ml per tube) and incubated for 10 minutes at 4°C. The cell suspension in each tube was diluted up to 2 mL with MACS buffer and centrifuged for 30 sec at 9300 g, 4°C. The supernatants were discarded and the cell pellets were resuspended in 1.5 ml MACS buffer per tube. The cell suspensions from each tube were transferred to two pre-wet LD columns (with MACS buffer, two LD columns for one brain sample, Miltenyi Biotec #130-042-901) and the cell flow-through were collected in 50 mL tubes on ice in a big covered Styrofoam cooler. The LD columns were rinsed twice with 2 mL MACS buffer. The flow-throughs were divided into multiple 2 mL tubes and centrifuged for 30 seconds at 9300 g, 4°C. The supernatants were discarded and the cell pellets were resuspended collectively in 1mL PBS for each sample. The myeloid cells were enriched by MACS using CD11b MicroBeads (Miltenyi Biotec # 130-049-601) according to manufacturer’s instructions. The cell viability was indicated by Trypan Blue and live/dead cell numbers were determined using an automated cell counter. Samples with cell viability more than 80% were subjected to single-cell RNA sequencing.

### Single-cell library preparation and RNA-sequencing

MACS-enriched brain myeloid cells were subjected to single-cell library preparation. For each sample approximately 12,000 cells were washed and resuspended in PBS containing 3% FBS and immediately processed as follows. Single-cell capture, barcoding and library preparation were performed using the 10X Chromium platform (10X Genomics), using version 3 chemistry according to the manufacturer’s protocol (10X Genomics #CG00052). The resulting cDNA and indexed libraries were checked for quality on an Agilent 4200 TapeStation, quantified by KAPA qPCR, and pooled for sequencing on 16.67% of lane of an Illumina NovaSeq 6000 S2 flow cell, targeting 6,000 barcoded cells with an average sequencing depth of 50,000 reads per cell. Illumina base call (bcl) files for the samples were converted to FASTQ files using CellRanger bcl2fastq (version 2.20.0.422, Illumina).

### Gene expression quantification from scRNA-seq data

The analysis pipeline of scBASE^9^ was used in order to avoid alignment bias due to differences in genetic background of mouse strains. First, we built the read alignment index by combining the custom strain-specific transcriptomes of CAST/EiJ, PWK/PhJ, WSB/EiJ, and C57BL/6J, created with g2gtools (http://churchill-lab.github.io/g2gtools). We removed PCR duplicates from the raw scRNA-seq data, and then aligned the remaining reads to the pooled transcriptome of the four strains using bowtie^47^ with ‘— all’, ‘—best’, and ‘—strata’ options. We processed the resulting bam files into an alignment incidence matrix (emase format) using alntools (https://churchill-lab.github.io/alntools) and quantified gene expression for each cell with emase-zero^10^ (https://github.com/churchill-lab/emase-zero). We collated the estimated UMI counts into a loom formatted file (http://loompy.org) for downstream analysis. A docker container in which all the above-mentioned software tools are pre-installed is freely available at https://hub.docker.com/r/kbchoi/asesuite-sc.

### Identification of brain myeloid cell types and microglia subtypes

First, we identified myeloid cell types by filtering out non-myeloid cells using a standard Seurat (v3.0)^11,48^ clustering pipeline for each strain, respectively. Cells with fewer than 600 detected genes or higher than 8% of mitochondrial genes were removed before initial analysis. We performed dimension reduction using PCA followed by UMAP using 3,000 most variable genes after normalizing the UMI counts. We identified marker genes for each clusters (FindAllMarkers) with default setting, and then annotated each cluster using enrichCellMarkers package^49^. We repeated the same clustering analysis after filtering out non-myeloid cells to refine PCA projection of myeloid cell types for each strain. We integrated myeloid cell clusters across the mouse strains (IntegrateData) and repeated the same clustering analysis. We identified a total of 91,201 myeloid cells including microglia, perivascular macrophage, monocytes and neutrophil (22,212 from B6, 24,976 from CAST, 20,192 from PWK and 23,821 from WSB, Fig. 1d, Supplementary Data Fig 1). Next, for microglia sub-clustering, we selected only those cells defined as microglia (unintegrated data) for integration and repeated the same clustering analysis. We identified a total of 87,746 microglia composed of 13 putative microglia subtypes (20,732 from B6, 24,124 from CAST, 19,702 from PWK and 23,188 from WSB, Fig. 3a, Supplementary Data Fig 1).

### Differential composition analysis of microglia subtypes

The proportion of microglia subtypes in each sample was calculated by dividing the number of cells in a given cluster by the total number of cells from each sample. A two-way ANOVA with Tukey’s post hoc test (aov and TukeyHSD function in base R) was employed to assess the strain, genotype, and strain by genotype effect on the percent of cells per cluster. Significance for genotype comparisons within strains were reported in each figure. The complete comparisons with confidence interval and adjusted p values (p.adj) are reported in Supplementary table 3.

### Differential gene expression, marker gene identification and correlation analysis

The strain, genotype and strain by genotype effect on single-cell gene expression for each microglia cluster was assessed by edgeR package^50^ (https://osca.bioconductor.org/). The single-cell microglia gene raw counts from a given cluster of each sample was summed as pseudo-bulk gene expression data before passing to standard DE gene analysis pipeline of edgeR using a quasi-likelihood method (glmQLFTest function). The gene expression model is built to access the strain, genotype and strain by genotype effect while regressing out batch effect (psedo-bulk gene expression/cluster ∼ strain + genotype + stran:genotype + batch). The complete DE gene analysis results with all coefficients for each cluster were reported in Supplementary Table 4 (FDR < 0.05 is considered significant). The initial myeloid and microglia marker genes for each clusters were determined using FindAllMarkers with the default Wilcoxon rank sum test in Seurat package, comparing gene expression of a given cluster to the rest of the clusters with all groups combined (Fig.2-4, p.adj < 0.05 was considered significant). For strain-specific microglia marker gene comparison (Fig. 6-7, Extended Data Fig 7), DAM, IRM, Ccl3/Ccl4-enriched microglia were compared to homeostatic microglia in each strain (genotype combined) using FindMarkers with Wilcoxon rank sum test in Seurat package. To estimate the similarity of microglia between strains, a spearman correlation was performed based on the fold change of top marker genes (|logFC| > 0.5, FDR < 0.05) of DAM, IRM, Ccl3/Ccl4-enriched microglia from each strain. To estimate the similarity of DAM of B6, CAST, PWK and WSB to those defined by Ido Amit’s group (Table S2 in their publication^3^), the spearman correlation was performed based on the fold change of top marker genes (|logFC| > 0.5, FDR < 0.05) of DAM (Extended Data Fig. 4b).

### Pseudotime analysis

We performed pseudotime analysis for microglia using ‘destiny’ package^51^, a diffusion map based-pseudotime inference. Because ‘destiny’ cannot generate a diffusion map of all 87,746 cells, we randomly sampled 1,000 cells from each group (8000 cells for 8 groups).The first 30 principal components from these cells were processed through the ‘dpt’ function to generate a diffusion map. The first dimension of the diffusion map was used as pseudotime axis. A histogram displaying the distribution of 1000 microglia of each group along the pseudotime were plotted, with microglia cluster color coded.

### Ingenuity Pathway Analysis (IPA)

DE genes comparing wild-derived strains to B6 for homeostatic microglia (cluster H: 0-5 combined), DAM (cluster 6), IRM (cluster 7) and Ccl3/Ccl4-enriched microglia (cluster 7) were subjected to Diseases and Functions (DF) and Regulatory Effect (RE) analysis of IPA. The DE genes uploaded to IPA was defined as FDR < 0.05 and |FC| > 2 for homeostatic microglia, |FC| > 1 for DAM, IRM, and Ccl3/Ccl4-enriched microglia. The higher FC threshold set for homeostatic microglia was because there are many more DE genes compared to other clusters when FC be set at 1 which is not computationally efficient for IPA. The top 20 (approximately) most significant terms of DF from any of comparisons (CAST vs B6, PWK vs B6, WSB vs B6) were visualized in a heatmap.

### Human AD GWAS gene selection

The human AD GWAS genes were selected from two recent meta-analyses (Table 1^31^ and Table S2^32^). The GWAS genes from both tables were combined and were overlapped with equivalent mouse genes in scRNA-seq data from this study. The GWAS genes were summarized in Supplementary Table 6.

## Author Contributions

HSY, KDO and GRH designed the study. HSY developed experimental protocols, prepared samples for single-cell RNA sequencing (scRNA-seq) and performed bioinformatics analysis. KDO and KJK generated and maintained mice for this study. KC built the pipeline for the quantification of gene expression for the scRNA-seq data. KC, DAS, GWC provided advice on the scRNA-seq data analysis. KDO, HSY, and GRH wrote the manuscript. All authors approved the final version.

## Data Availability

All mouse strains are available through The Jackson Laboratory and all associated data are being made available through the Accelerating Medicines Partnership-Alzheimer’s disease (AMP-AD) knowledge portal. All analysis scripts are being made available through github.

## Acknowledgements

We gratefully acknowledge the contribution of Sandy Daigle, Michael Samuels and Paul Robson at the Single Cell Biology Laboratory at The Jackson Laboratory (JAX) for expert assistance with this publication. We thank Duy Pham from the Churchill lab at JAX for advice on pre-processing of scRNA-seq data on high-performance computing cluster. We thank Christoph Preuss from the Carter lab at JAX for advice on human GWAS AD gene analysis. We thank Sandeep Namburi at JAX research IT for installing essential packages and troubleshooting on RStudio server.

## Competing interests

The authors declare no competing interests.

## Funding Sources

This study was supported in part by the National Institute of Aging RF1AG051496 (GRH, HSY, KDO), RF1AG055104 (GRH, KDO), and Alzheimer’s Association Research Fellowship 2018-AARF-589154 (HSY).

## Figure Captions

**Extended Data Fig. 1 | Quality control for single-cell RNA-seq. a-d**, Box/violin plots showing the distribution of the number of total RNA reads (nCount_RNA), the number of total gene detected (nFeature_RNA) and the percent mitochondrial genes in all individual cells after each quality control steps. **a**, All recovered cells after gene expression quantification by emaze-zero. **b**, Remaining cells after removing cells with high-percent mitochondrial genes (>8%). **c**, Remaining cells after myeloid cell integration based on strain. **d**, Remaining cells after microglia integration based on strain.

**Extended Data Fig. 2 | Quality control for single-cell RNA-seq featuring on immediate early genes in microglia. a-b**, UMAP plots (a) and dot plot (b) showing the expression of several common immediate early genes including Fos, Fosb, Dusp1, Nr4a1, Arc and Egr1 in integrated microglia clusters (Fig. 3a).

**Extended Data Fig. 3 | Marker genes in homeostatic microglia and variation in the percent of microglia subtype. a**, Dot plot showing the expression of top microglia marker genes of all clusters (Fig. 3b) in homeostatic microglia subtypes (clusters 0, 1, 2, 3, 4, 5) and hyper-homeostatic microglia (cluster 8). **b**, Dot plot showing the expression of the top microglia marker genes of in homeostatic microglia (clusters 0,1, 2, 3, 4, 5) and 8. **c**, UMAP plot showing microglia clusters in each strain and genotype. **d**, The percent of 8 microglia subtypes in each replicate of each strain and genotype.

**Extended Data Fig. 4 | Comparisons of Disease Associated Microglia (DAM). a**, Volcano plot showing the top up- and down-regulated genes comparing DAM cluster 12 to DAM cluster 6. Cluster 12 showed increased expression in ribosomal genes and Tyrobp (log2FC > 0) and decreased expression in homeostatic microglia genes including Cx3cr1, Tgfbr1, Csf1r and Tgfbr2 (log2FC < 0). **b**, Spearman correlation coefficient matrix comparing top DAM (cluster 6) marker genes between B6, CAST, PWK, WSB and Ido Amit’s DAM gene based on fold change (log2FC > 0.5, FDR < 0.05).

**Extended Data Fig. 5 | Features of number of total genes and the percent of mitochondrial and ribosomal genes in all microglia subtypes**. Box/violin plots showing the distribution of the number of total gene detected (nFeature_RNA), the percent of mitochondrial (percent.mt) and ribosomal (percent.ribo) genes in cells from all identified microglia subtypes. * adjusted p value (p.adj) < 0.05. All comparisons were made by comparing the medians in clusters 6, 7, 8, 9, 10, 11 and 12 to combined homeostatic microglia (cluster 0 to 5), respectively, using one-way ANOVA followed by Tuskey’s post hoc test. The complete cross-cluster comparison results were shown in Supplementary Table 2.

**Extended Data Fig. 6 | Marker genes and cell abundance comparison in microglia subtypes 9, 10 and 11. a-b**, UMAP and violin plots highlighting marker genes in Ccl3/Ccl4-enriched (cluster 10) and proliferative (cluster 11) microglia. **c-e**, Box plots showing the percent of ribosomal gene-enriched microglia (cluster 9), cluster 10 and 11 microglia in all group of mice. Strain, genotype and strain by genotype effects were assessed by 2-way ANOVA followed by Tukey’s post hoc test. All comparisons (comparing WT and *APP/PS1* within each strain for a given cluster) are significant (adjusted p value, p.adj < 0.05) except for those labelled with NS (not significant, p.adj ≥ 0.05). Detailed p.adj values and confidence intervals for within and cross strain/genotype comparisons were reported in Supplementary Table 3.

**Extended Data Fig. 7 | Strain-specific gene expression in Ccl3/Ccl4-enriched microglia (cluster 10). a**, Diagram illustrating the strain-specific DE gene analyses in Ccl3/Ccl4-enriched microglia (cluster 10). IRM marker genes comparing cluster 10 to homeostatic microglia (FDR < 0.05) of each strain were subjected to venn diagram, spearman correlation and scatter plots. DE genes comparing wild-derived IRM to B6 IRM were subjected to volcano plot and IPA. **b**, Venn diagram showing the overlap of the numbers of cluster 10 marker genes across strains. **c**. Spearman correlation matrix based on FC of top IRM marker genes (|log2FC| > 0.5) between strains showing the correlation coefficients. All correlations are significant (p < 0.05). **d-f**, Scatter plots showing IRM marker genes between CAST vs B6 (d), PWK vs B6 (e) and WSB vs B6 (f). B6 (dark grey): significant only in B6 but not in wild-derived strains; CAST (green), PWK (red), WSB (purple): significant only in CAST, PWK or WSB but not in B6; Both (yellow): significant only in both wild-derived and B6 mice. **g-i**, Volcano plots showing DE genes comparing CAST vs B6 (g), PWK vs B6 (h) and WSB vs B6 (i). Top 25 gene DE genes based on FC and FDR were labelled. **j**, Heat map summarizing top 21 significantly enriched terms of Diseases and Functions based on DE genes from comparisons of wild-derived vs B6 mice (pval-BH < 0.05, |z-score| ≥ 2). **k**, Selected Regulatory Effects for CAST vs B6 highlighting network of ‘Mobilization of lymphatic system cells’, ‘Extravasation of cells’ and ‘Chemoattraction of monocytes’. The color code is the same as described in Fig.5 g-h.

**Supplementary Table 1 | Microglia cluster marker gene list with all strains and genotypes combined.** Marker genes are resulted from FindAllmarkers function in Seurat package with the parameter set as min.pct = 0.2, logfc.threshold = 0.25, max.cells.per.ident=500.

**Supplementary Table 2 | One-way ANOVA followed by Tukey post hoc testing on number of feature gene (nFeature_RNA), percent of mitochondria genes (percent_mito), percent of ribosomal genes (percent_ribo) between microglia subclusters.** The significance threshold was set at the adjusted p value (p.adj) less than 0.05. “diff” stands for the difference of mean for a given comparison. “lwr” and “upr” stands for lower and upper range of confidence interval at the significance level of 0.05.

**Supplementary Table 3 | Two-way ANOVA followed by Tukey post hoc testing on the percent of cells between clusters as strain, genotype and strain by genotype effect.** Three sets of comparisons (*APP/PS1* vs WT within strain, WT between strains, *APP/PS1* between strains) were listed in individual sheets. The significance threshold was set at the adjusted p value (p.adj) less than 0.05. “diff” stands for the difference of mean for a given comparison. “lwr” and “upr” stands for lower and upper range of confidence interval at the significance level of 0.05.

**Supplementary Table 4 | Strain, genotype and strain by genotype gene list for each cluster (H, 6, 7, 8, 9, 10, 11, 12)** The specific effect is listed as “coef” column and detailed below. logFC is 2 based. We recommend using Excel data filter function or R to query genes based on their coef and cluster. **StrainCAST**: strain effect comparing CAST vs B6, positive logFC indicates gene expression is higher in CAST than B6, and vice versa. **StrainPWK**: strain effect comparing PWK vs B6, positive logFC indicates gene expression is higher in PWK than B6, and vice versa. **StrainWSB**: strain effect comparing WSB vs B6, positive logFC indicates gene expression is higher in WSB than B6, and vice versa. **StrainCAST-GenotypeAPP-PS1**: strain by genotype effect in CAST, positive logFC indicates gene expression is higher in CAST than B6 in response to *APP/PS1*, and vice versa. **StrainPWK-GenotypeAPP-PS1**: strain by genotype effect in PWK, positive logFC indicates gene expression is higher in PWK than B6 in response to *APP/PS1*, and vice versa. **StrainWSB-GenotypeAPP-PS1**: strain by genotype effect in WSB, positive logFC indicates gene expression is higher in WSB than B6 in response to *APP/PS1*, and vice versa. **GenotypeAPP-PS1**: genotype effect comparing *APP/PS1* vs WT, positive logFC indicates gene expression is higher in *APP/PS1* than B6.

**Supplementary Table 5 | Marker genes for DAM, IRM and *Ccl3*/*Ccl4*-enriched microglia for each strain.**

**Supplementary Table 6 | Comparison of human AD GWAS genes with mouse strain-effect genes in scRNA-seq dataset.**

## Notes

### Competing Interest Statement

The authors have declared no competing interest.

