## Supplementary figures and images for "Natural genetic variation determines microglia heterogeneity in wild-derived mouse models of Alzheimer’s disease"

### Extended Data Figure 1

Extended Data Fig. 1

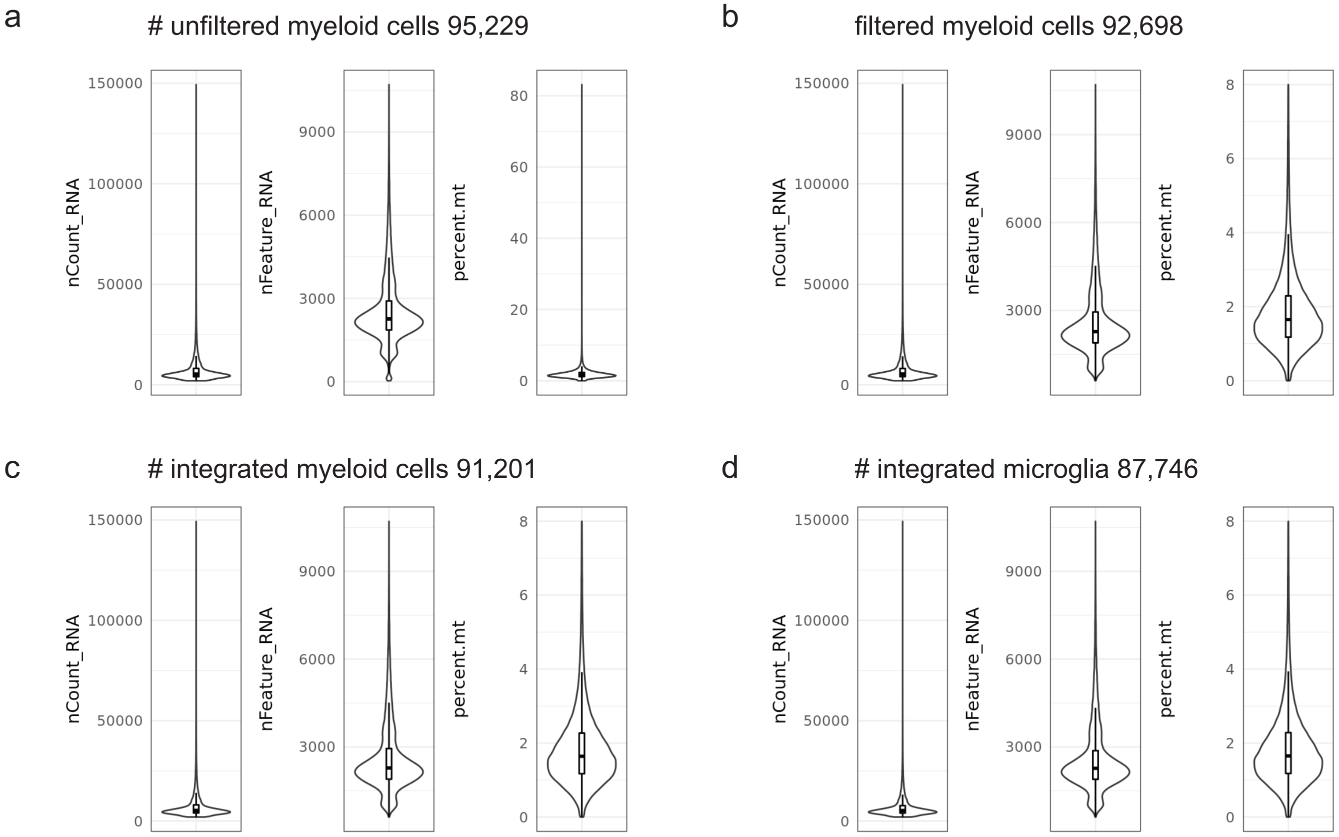

### Extended Data Figure 2

Extended Data Fig. 2

a

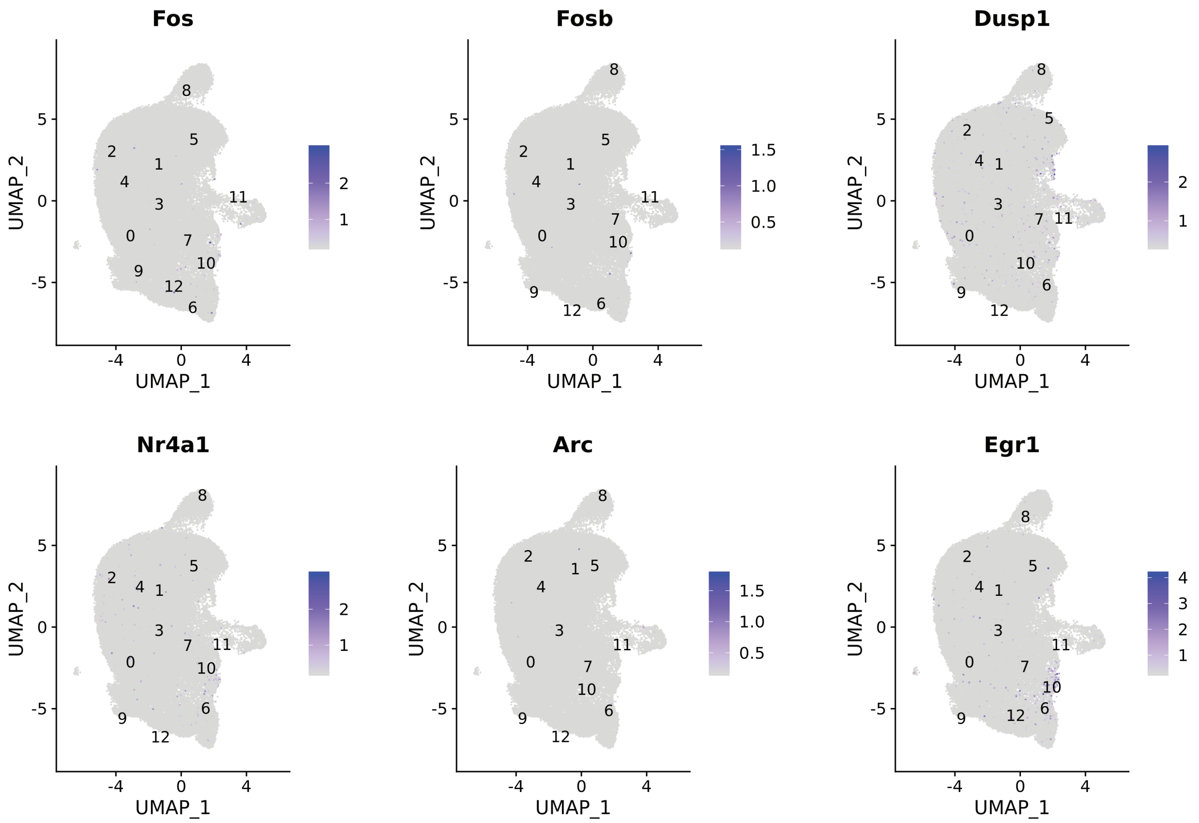


b

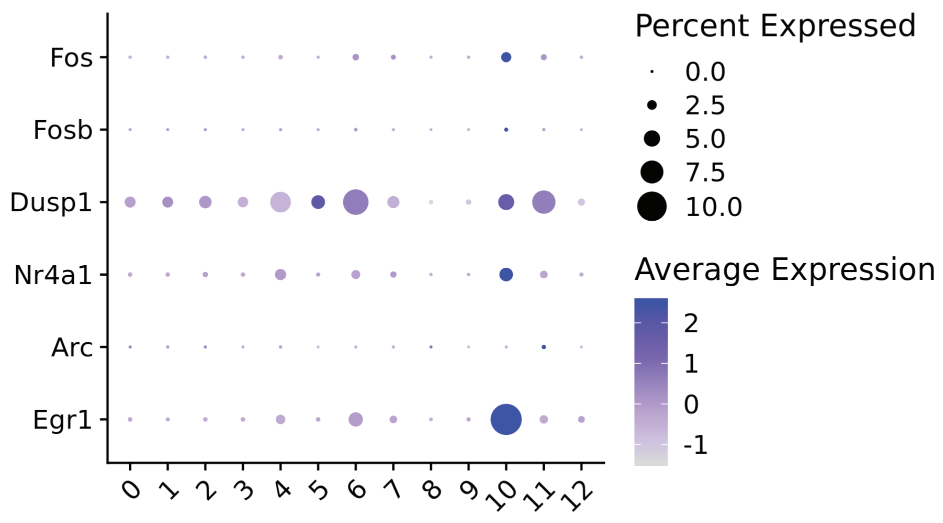

### Extended Data Figure 3

**Extended Data Fig. 3**

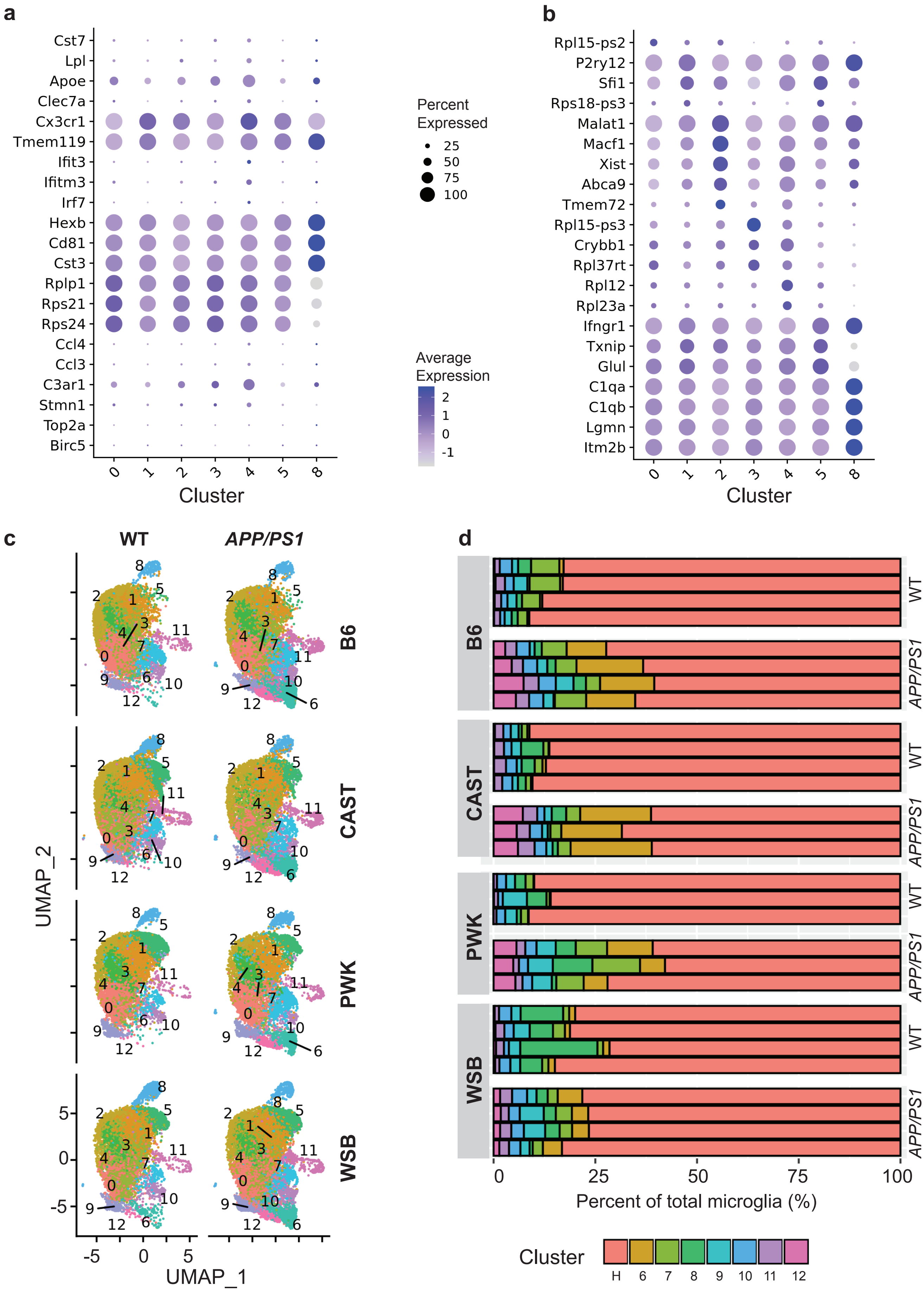

### Extended Data Figure 4

**Extended Data Fig.4**

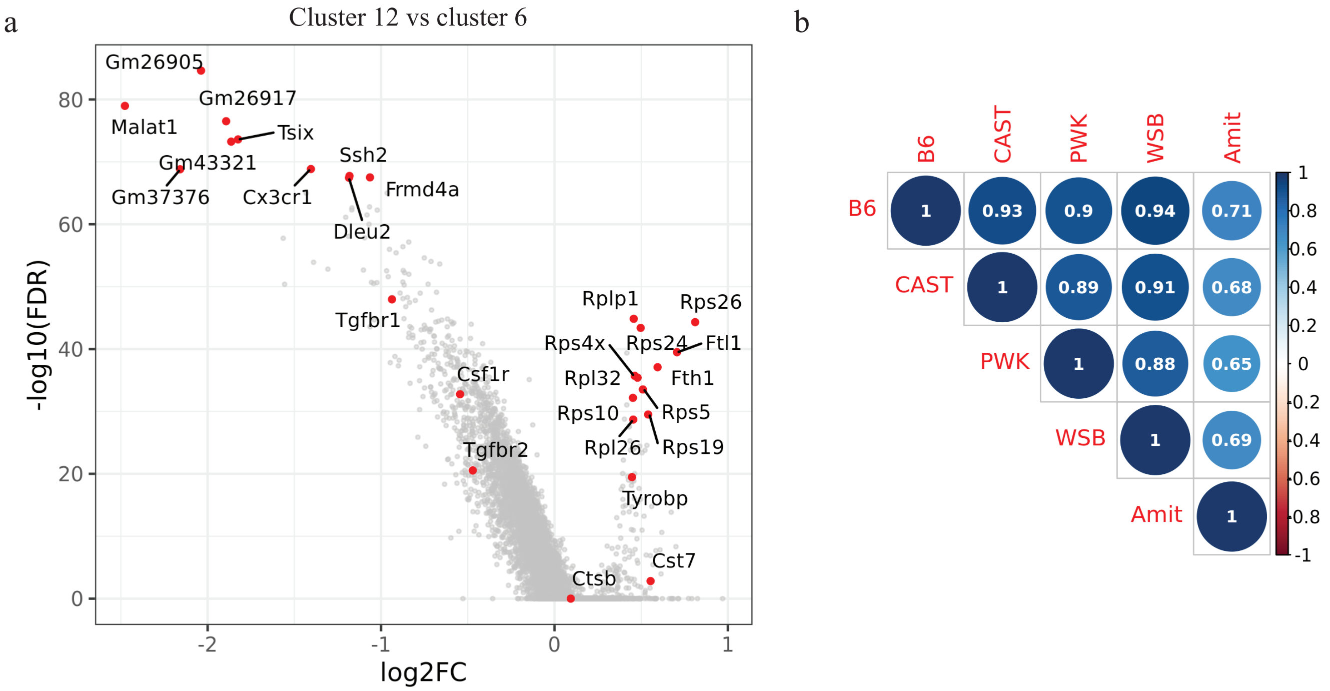

### Extended Data Figure 5

Extended Data Fig.5

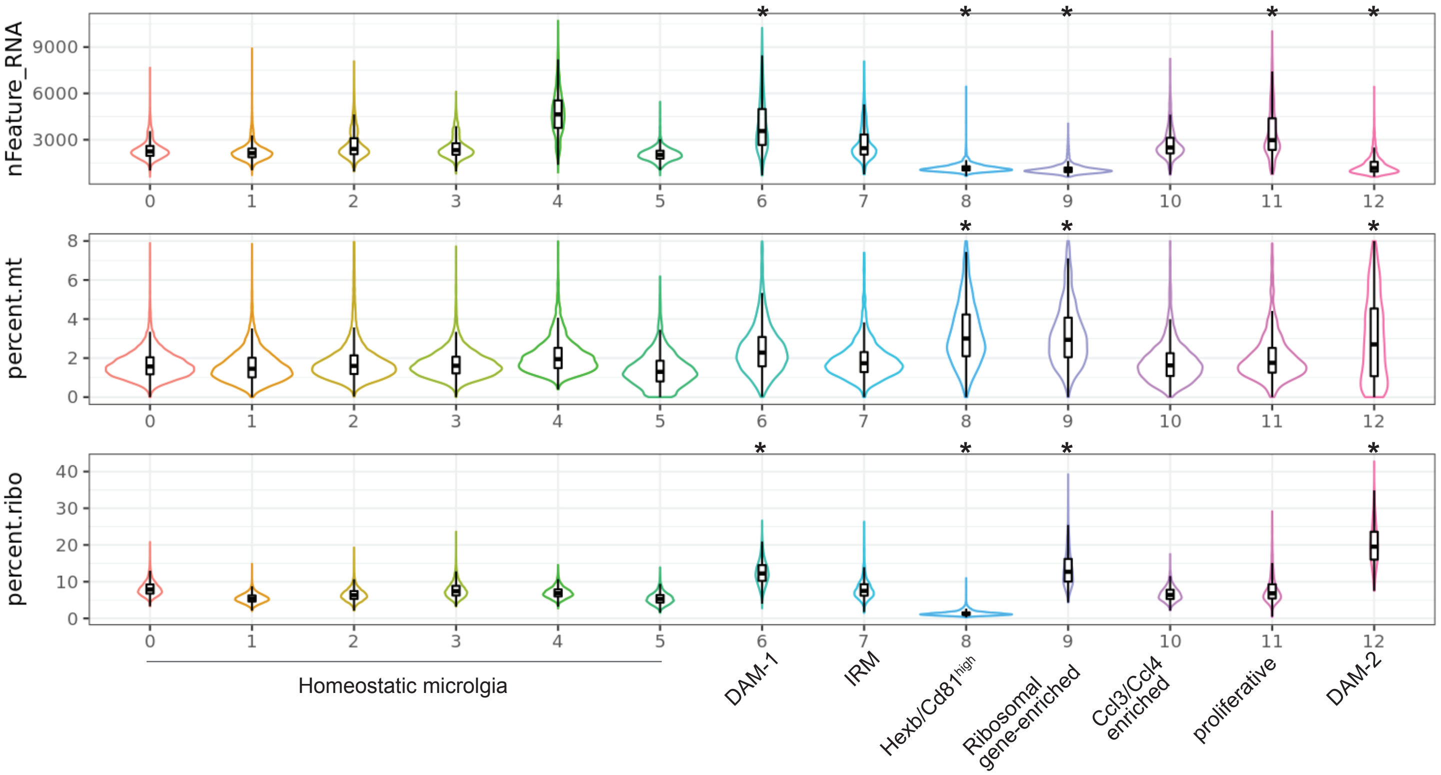

### Extended Data Figure 6

#### Extended Data Fig.6

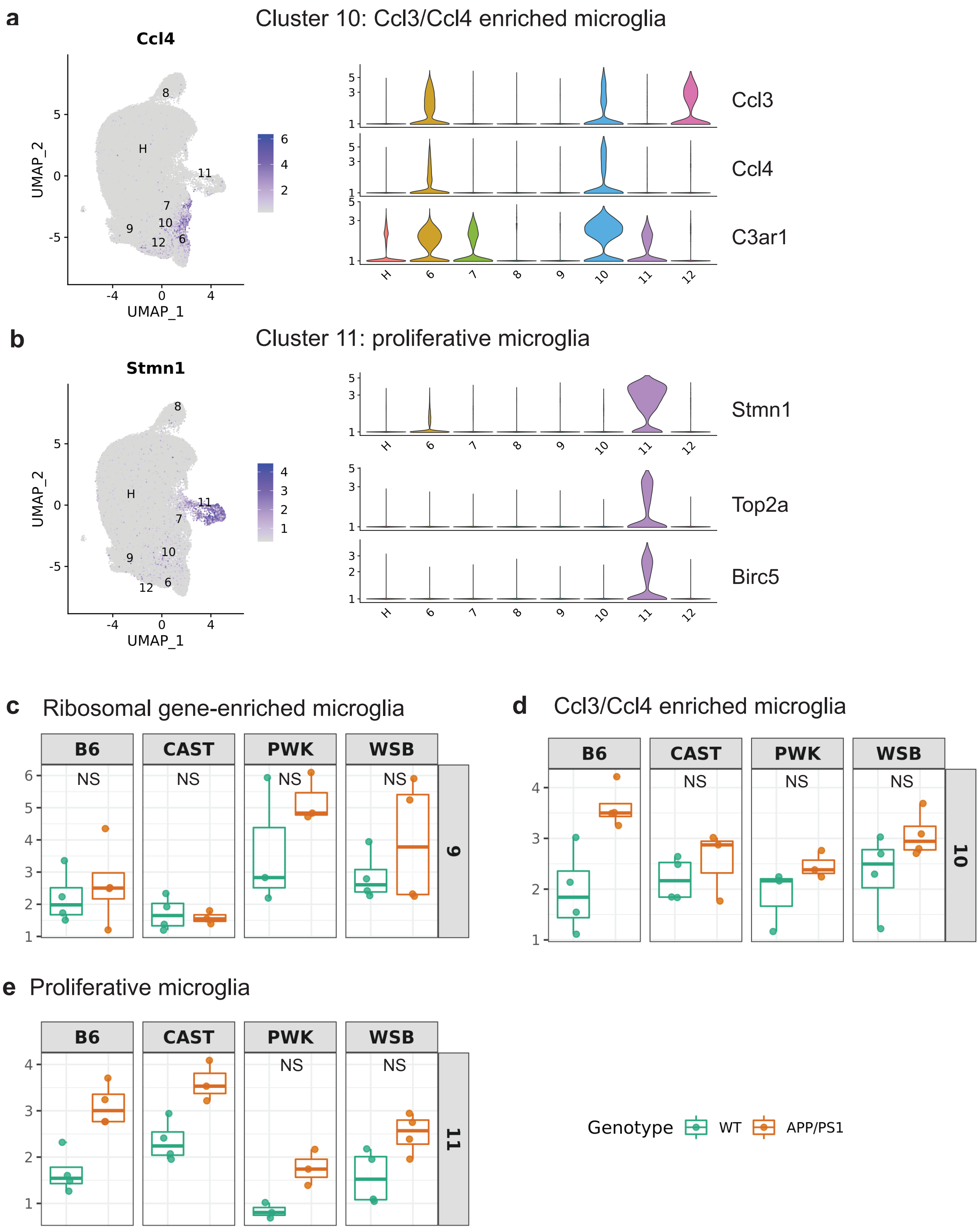

### Extended Data Figure 7

Extended Data Fig. 7

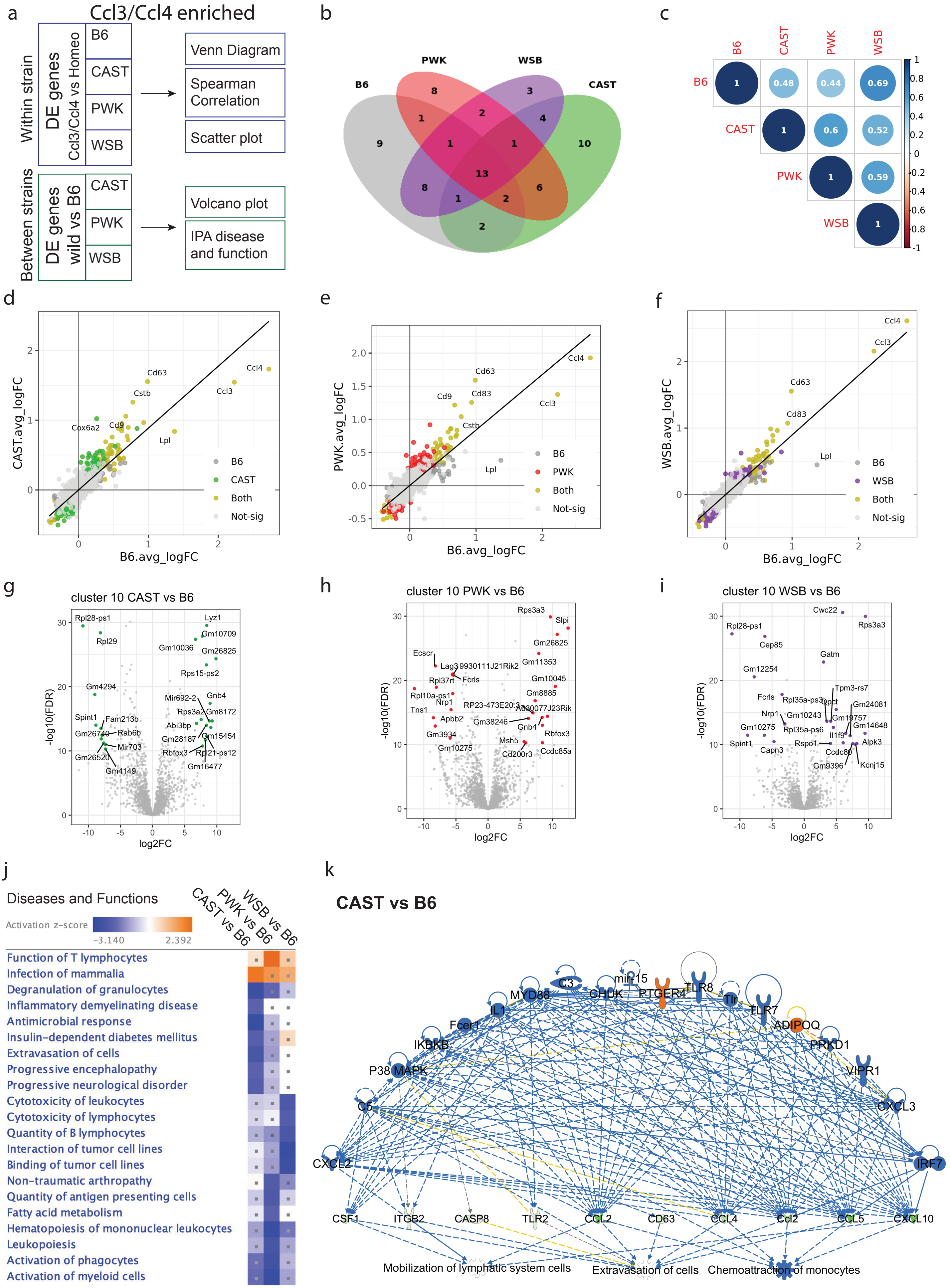
